# Myeloid-specific KDM6B inhibition sensitizes Glioblastoma to PD1 blockade

**DOI:** 10.1101/2022.11.28.518243

**Authors:** Sangeeta Goswami, Deblina Raychaudhuri, Pratishtha Singh, Seanu Meena Natarajan, Yulong Chen, Candice Poon, Mercedes Hennessey, Jan Zhang, Swetha Anandhan, Brittany Parker Kerrigan, Marc D. Macaluso, Zhong He, Sonali Jindal, Frederick F. Lang, Sreyashi Basu, Padmanee Sharma

## Abstract

Glioblastoma (GBM) tumors are enriched in immune-suppressive myeloid cells and are refractory to immune checkpoint therapy (ICT). Targeting epigenetic pathways to reprogram the functional phenotype of immune-suppressive myeloid cells to overcome resistance to ICT remains unexplored. Single-cell and spatial transcriptomic analyses of human GBM tumors demonstrated high expression of an epigenetic enzyme - histone 3 lysine 27 demethylase (KDM6B) in intra- tumoral immune-suppressive myeloid cell subsets. Importantly, myeloid-cell specific *Kdm6b* deletion enhanced pro-inflammatory pathways and improved survival in GBM tumor-bearing mice. Mechanistic studies elucidated that the absence of *Kdm6b* enhances antigen-presentation, interferon response and phagocytosis in myeloid cells by inhibiting mediators of immune suppression including *Mafb, Socs3* and *Sirpa*. Further, pharmacological inhibition of KDM6B mirrored the functional phenotype of *Kdm6b* deleted myeloid cells and enhanced anti-PD1 efficacy. Thus, this study identified KDM6B as an epigenetic regulator of the functional phenotype of myeloid cell subsets and a potential therapeutic target to improve response to ICT.

## Main

Immune cells of the myeloid lineage constitute a dominant portion of the tumor immune microenvironment and demonstrate significant plasticity depending on cues received from the environment ^1, 2^. While pro-inflammatory myeloid cells are crucial for mounting an effective anti- tumor immune response, immune-suppressive myeloid cells are associated with poor prognosis and therapeutic resistance in multiple cancer types ^3–6^. Emerging evidence suggests that immune- suppressive myeloid cells play a critical role in both primary and adaptive resistance to immunotherapy ^7–11^. Therefore, developing therapeutic strategies to target immune-suppressive myeloid cells is a critical approach to enhance response to cancer immunotherapy and has been the focus of intense research for many years ^12–14^ .

Most studies have focussed on depleting intratumoral immune-suppressive myeloid cells, blocking their trafficking, and targeting individual immune-suppressive pathways to enhance anti- tumor immunity ^15–20^. However, many of these strategies have not been successfully translated to the clinic partly due to the functional heterogeneity and redundancy of pathways in myeloid cell subsets. Newer technologies such as single cell RNA sequencing (scRNA seq) further demonstrated the wide spectrum of functional states attained by each of these subsets based on signals received from the niche they inhabit ^21–25^. This significant plasticity of myeloid cells highlights the important role of epigenetic regulation of their cell state ^26, 27^. However, the impact of epigenetic regulation of intratumoral myeloid cell plasticity on therapeutic resistance remains largely unexplored.

Glioblastoma (GBM) is an aggressive form of brain tumor, highly infiltrated with immune- suppressive myeloid cells and demonstrates resistance to ICT ^28–31^. We have previously shown the persistence of tumor-associated macrophages (TAMs) in the GBM tumor microenvironment even after treatment with anti-PD1 therapy ^32^. In this study, we aimed to identify epigenetic factors regulating the functional phenotype of intratumoral myeloid cell subsets in order to reprogram these cells to a pro-inflammatory state thus enhancing anti-tumor immunity and efficacy of ICT. scRNA seq of GBM tumors resected from patients demonstrated high expression of histone 3 lysine 27 demethylase (KDM6B) in myeloid cell subsets including monocytes, macrophages and dendritic cells (DCs). Further, spatial transcriptomic analysis of human GBM tumors showed significant infiltration of *KDM6B* expressing immune-suppressive myeloid cells in the tumor microenvironment (TME). KDM6B is an epigenetic enzyme that demethylates the repressive trimethylation mark at histone 3 lysine 27 (H3K27me3) thereby promoting gene transcription ^33^. Importantly, in murine models of GBM, LysM^cre^KDM6B^fl/fl^ micecarrying *Kdm6b* deletion in myeloid cells had enhanced pro-inflammatory pathways and improved survival compared with their wild-type counterparts. Single-cell assay for transposase-accessible chromatin sequencing (scATAC seq) and chromatin immunoprecipitation followed by sequencing (ChIP seq) demonstrated that KDM6B directly regulates H3K27me3 enrichment of genes including *Mafb, Socs3* and *Sirpa* which inhibit critical pro-inflammatory pathways such as cytokine production and phagocytosis in macrophages^19, 20, 34, 35^, providing mechanistic insight into enhanced pro-inflammatory pathways noted in the absence of *Kdm6b*. Further, pharmacological inhibition of KDM6B could recapitulate the functional phenotype of *Kdm6b* deleted myeloid cells and improve sensitivity to anti-PD1 therapy in GBM. Together, these findings have provided critical insight into KDM6B-mediated epigenetic regulation of intratumoral myeloid cell functions. Overall, this study proposes a new paradigm of targeting the epigenetic machinery to regulate intratumoral myeloid cell plasticity thus reprogramming them into a pro-inflammatory phenotype to overcome myeloid cell-mediated resistance to ICT.

### Single-cell and spatial transcriptomic analyses of human GBM tumors demonstrated selective expression of *KDM6B* in immune-suppressive myeloid cells

We performed scRNA seq of intratumoral *CD45+* cells in GBM tumors resected from patients (n=5). The patient characteristics are highlighted in Supplementary Table 1. Unsupervised clustering and uniform manifold approximation and projection (UMAP) analyses revealed distinct *CD3E+CD4+*T cells (C2), *CD3E+CD4+FOXP3*+ regulatory T cells (C14), *CD3E+CD8A+*T cel (C5,7,9,10,11), *KLRK1+*NK cell (C13)*, CD19+CD79+* B cell (C15), and myeloid cell subsets (C0,1,4,3,6,8,16) (Fig.1A, Extended Data Fig. 1A). Next, to investigate the intratumoral myeloid cells in human GBM samples in greater detail, we re-clustered myeloid cell populations characterized by the expression of *CD68* (Fig. 1B). Annotation of each of these individual clusters (Fig. 1B-C) revealed the presence of three microglial-like clusters (C0,4 and 8), one *STAB1+LYVE1+DAB2+* brain-associated macrophage cluster (C3), a *CHI3L1+TIMP3+SPARC+COL1/3/4A1+* population (C9), a *CD1C+CD1A+AREG+* dendritic cell cluster (C6) and four distinct monocytic/macrophage populations (C1, C2, C5, and C7).

**Fig. 1:**
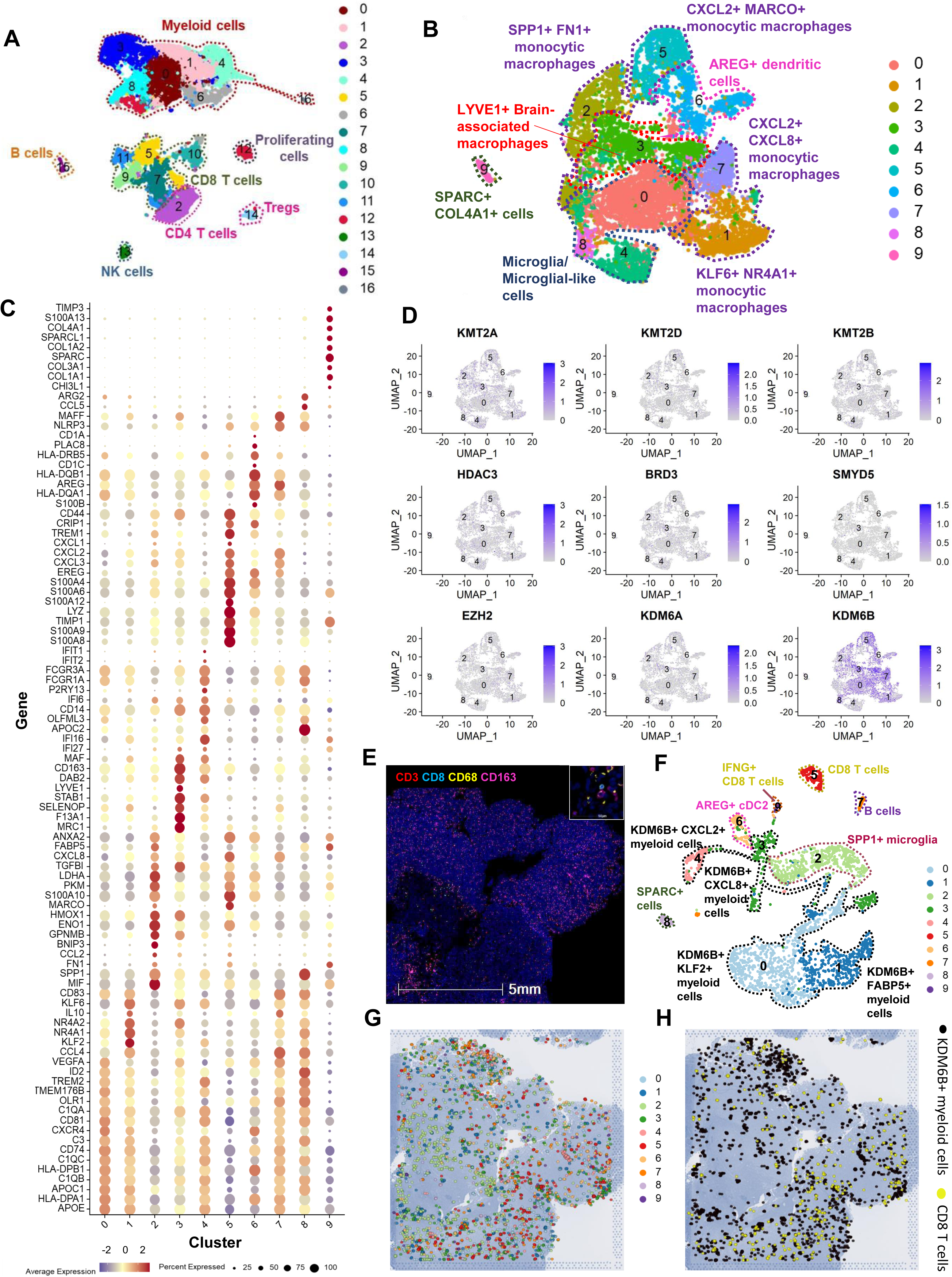
Single-cell and spatial transcriptomic analyses of human GBM tumors demonstrated selective expression of KDM6B in immune-suppressive myeloid cells. **(A)** UMAP plot of scRNA seq data depicting the different immune cell subsets (CD45+ cells) in the TME of patients with GBM (n=5). **(B)** UMAP plot of myeloid cells obtained by reclustering the CD68+ clusters from Fig. 1A. **(C)** Dotplot showing the average expression of indicated genes as well as the percentage of cells expressing the gene in the indicated myeloid clusters shown in Fig. 1C. **(D)** Feature plots of scRNA seq data demonstrating the expression of indicated genes encoding epigenetic enzymes, in intratumoral myeloid cell clusters. **(E)** Multiplex immunofluorescence staining shows distribution of CD3 (red), CD8 (cyan), CD68 (yellow) and CD163 (magenta) in a GBM case. Inset shows CD3+, CD8+ and CD68+CD163+ myeloid cells. **(F)** UMAP plot of scRNA seq data from patient 6 (Supplementary Table 1), showing CD45+ immune cell subsets in the GBM TME used to embed single cells to their spatial coordinates in tissue sections by applying CellTrek. **(G)** Gene expression data from all the different immune cell clusters from the matched patient plotted on spatial coordinates. **(H)** Gene expression data from the KDM6B expressing myeloid cell clusters (black) and CD8 T cell clusters (yellow), plotted on spatial coordinates.

Out of the three microglial clusters, C4 has several pro-inflammatory markers including interferon signature genes (ISGs)- *IFI27, IFI16, IFI6* (Fig. 1C). In contrast, C0 (*APOE+TREM2+CD81+OLR1+HLA-DPA+C1Q+)* and C8 (*OLFML3+ P2RY13+*) are noted to express suppressive markers such as *VEGFA, CCL4* (C0) and *SPP1, CCL4, ARG2* (C8) respectively (Fig. 1C). Expression of immune-suppressive markers such as *MRC1, CD163, TGFBI and SELENOP* was also noted in brain- associated macrophage cluster (C3) (Fig. 1C). Additionally, all the monocytic macrophage clusters display a predominantly suppressive phenotype. We noted the presence of a pro-angiogenic/hypoxic *FN1+SPP1+MIF+BNIP3+HMOX1+CXCL8+ANXA2+* cluster (C2), a *KLF2+KLF6+IL10+NR4A1+* cluster (C1), a *S100A9+MARCO+CXCL1+CXCL2+CXCL8+CRIP1+ANXA2+* cluster (C5) and a *CCL4+CXCL2+CXCL8+IL10+AREG+NLRP3+* cluster (C7) (Fig.1C). Overall, this analysis highlighted the enrichment of suppressive myeloid cell subsets in GBM tumors resected from patients.

To identify epigenetic factors which are critical for the regulation of immune-suppressive phenotype and function in the tumor microenvironment (TME), we investigated gene expression of several canonical epigenetic modifiers previously implicated in the regulation of myeloid cell polarization and function (Fig. 1D)^26, 27^. Interestingly, amongst the selected epigenetic enzymes, we noted high levels of expression and selective enrichment of *KDM6B/JMJD3* in the myeloid cells, specifically in the subsets expressing immune-suppressive markers such as *CSF1R*, *KLF2, KLF6, CXCL8 and SPP1* (Extended Data Fig.1B-C). H3K27 methylation is an important epigenetic determinant of myeloid cell phenotype and function ^36^. *KDM6B* works in tandem with other epigenetic modifiers such as *KDM6A* and *EZH2* in the regulation of H3K27 mediated gene expression . Unlike *KDM6B, KDM6A* and *EZH2* showed minimal expression in the intratumoral myeloid cell subsets (Fig.1D). Next, to confirm our findings, we used two independent scRNA seq datasets with n=4^32^ and n=20 ^37^ GBM patients respectively. Analyses of these two datasets demonstrated distinct *CD3E+* T cell and *CD68+* myeloid cell clusters (Extended Data Fig. 2A-D). Importantly, similar to our primary cohort of GBM patients, *KDM6B* enrichment was observed in intratumoral immune-suppressive myeloid cells (Extended Data Fig. 2A-D).

Next, we performed spatial transcriptomic analysis of GBM tumors (n=3) resected from patients (charcteristics enlisted in Supplementary Table 1) using the 10X-Genomics Visium platform. Hematoxylin and Eosin (H&E) staining was done to determine the overall architecture of each tumor section (Fig S3A). Immunofluorescence microscopy demonstrated the presence of intratumoral CD3+, CD8+ T cells and CD68+CD163+ myeloid cells (inset) and confirmed that human GBM tumors are diffusely infiltrated with myeloid cells (Fig. 1E). Further, immunohistochemical (IHC) staining confirmed the expression of KDM6B protein in all the human GBM sections studied (Extended Data Fig.3B). To visualize the spatial localization of *KDM6B*+ immune-suppressive myeloid cells in the GBM TME, we used matched scRNA seq data to embed single cells to their spatial coordinates in tissue sections by applying CellTrek ^38^. Multiple myeloid cell, T cell, B cell, and NK cell clusters could be spatially delineated based on characteristic gene signatures (Fig.1F,G, Extended Data Fig.4A-D and Extended Data Fig.5A-C). As expected, we noted inter-tumoral qualitative and quantitative heterogeneity in the spatial landscape of immune cell subsets. However, enrichment of *KDM6B* expressing myeloid clusters co-expressing immune-suppressive markers such as *SPP1, CXCL8, MARCO* and *MAFB* was observed across all the tumor sections studied (Fig. 1F-H, Extended Data Fig. 4A-F and Extended Data Fig. 5A-C).

Together, single-cell and spatial transcriptomic analyses of human GBM tumor samples demonstrated the expression of KDM6B in the immune-suppressive myeloid cell subsets in patients with GBM.

### Myeloid cell-specific deletion of *Kdm6b* improves survival in preclinical models of GBM and results in a pro-inflammatory milieu in murine GBM tumors

To interrogate the impact of KDM6B-mediated regulation of the functional phenotype of myeloid cell subsets on anti-tumor immunity, we generated a murine model bearing myeloid cell-specific deletion of the *Kdm6b* gene (LysM^cre^KDM6B^fl/fl^) (Extended Data Fig. 6A,B). Mass cytometry (CyTOF) based immunophenotyping studies showed no significant differences in the relative abundance of the major immune cell subsets including myeloid cells in the immune cell repertoire of bone marrow, spleen, and lymph-node from wild type (control) and LysM^cre^KDM6B^fl/fl^ mice (Extended Data Fig.6C), indicating that myeloid-specific deletion of *Kdm6b* does not alter the development of immune cell populations. Next, we orthotopically inoculated murine GBM cell line -GL261 into the brain of control and LysM^cre^KDM6B^fl/fl^ mice. Magnetic Resonance Imaging (MRI) studies done on day 14 post tumor inoculation revealed a lower tumor burden in mice having myeloid-cell specific *Kdm6b* deletion as compared with control (Fig. 2A). We also observed a similar reduction in tumor volumes in LysM^cre^KDM6B^fl/fl^ mice bearing another murine GBM cell line- CT-2A (Fig. 2A). In addition, survival studies showed improvement in survival of both GL261 and CT-2A GBM tumor-bearing LysM^cre^KDM6B^fl/fl^ mice compared with control (Fig. 2B). Thus, *Kdm6b* expression in myeloid cell subsets potentially aids in maintaining the suppressive phenotype as constitutive deletion of *Kdm6b* in myeloid cells attenuated tumor growth and provided a survival advantage in two pre-clinical models of GBM tumor bearing mice.

**Fig. 2:**
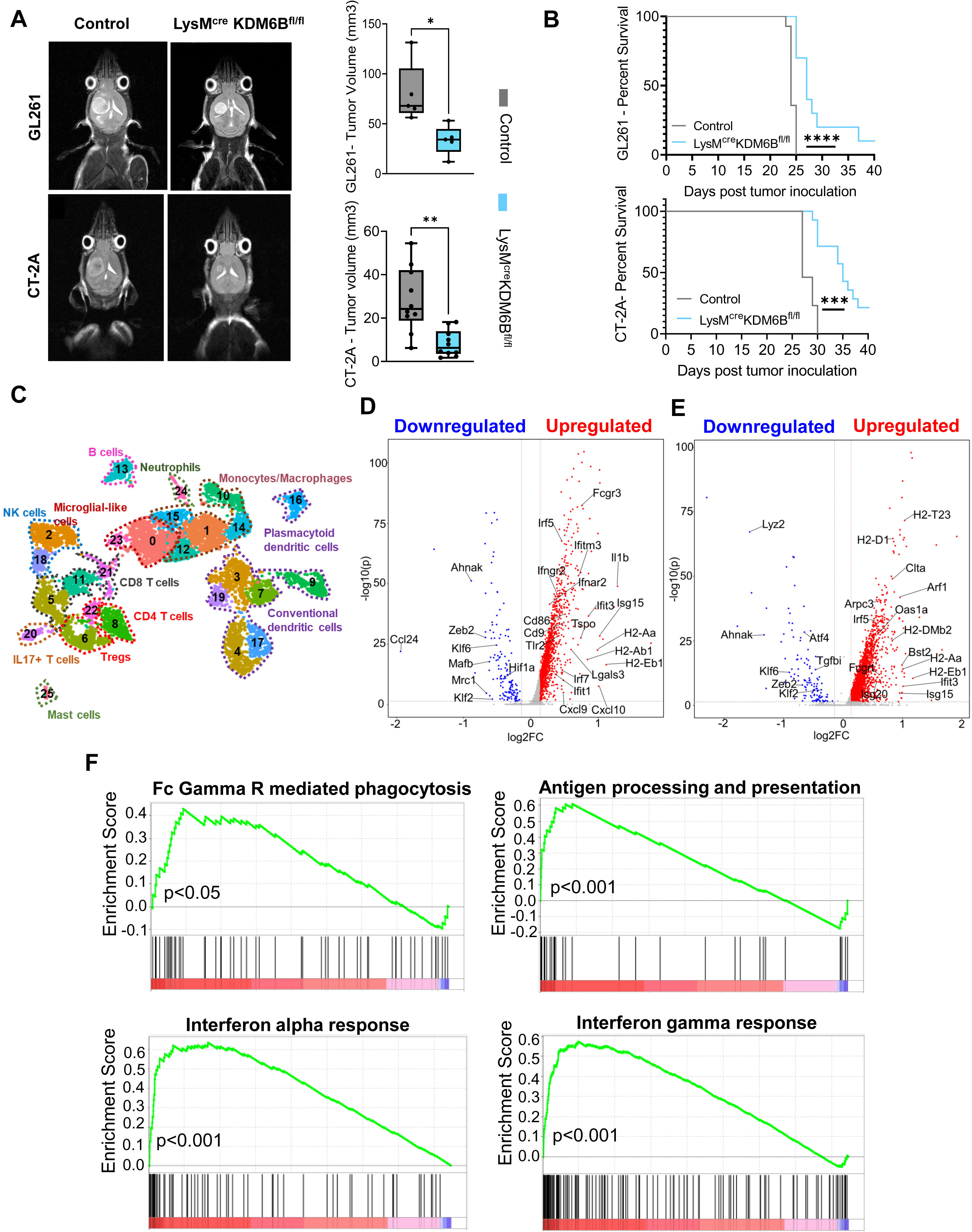
Myeloid cell-specific deletion of Kdm6b improves survival in preclinical models of GBM and results in a pro-inflammatory milieu in murine GBM tumors. **(A)** Representative axial MRI images taken on day 14 post tumor inoculation, of GL261 (top panel) and CT-2A (bottom panel) tumors in control (left panel) and LysM^cre^KDM6B^fl/fl^ mice (right panel). Box and whisker plot demonstrating the difference in tumor volume as calculated from MRI images of GL261 (top panel) (n=5/group) and CT-2A (bottom panel) (n=10/group) GBM tumors from control and LysM^cre^KDM6B^fl/fl^ mice. Two-tailed Student’s t-test was performed (* p<0.05, ** p<0.01). **(B)** Kaplan Meier plot depicting the difference in survival of GL261 (top panel) and CT-2A (bottom panel) tumor-bearing control and LysM^cre^KDM6B^fl/fl^ mice (n=10/group). Log-rank test was performed (***p<0.001, **** p<0.0001). Data is representative of 3 independent experiments. **(C)** UMAP plot of scRNA seq data showing the different immune cell subsets (Cd45+ cells) in the GBM (GL261) TME of control and LysM^cre^KDM6B^fl/fl^ mice (pooled 4-5 samples/group). The data is representative of 2 independent scRNA seq experiments. **(D)** Volcano plot representing differentially expressed genes between control and LysM^cre^KDM6B^fl/fl^ mice in intratumoral monocytes, macrophages and microglial-like cells (C0,1,10,12,14,15,23 in Fig. 2C). **(E)** Volcano plot depicting differentially expressed genes between control and LysM^cre^KDM6B^fl/fl^ mice in intratumoral DCs (C3,4,7,9,16,17,19 in Fig. 2C). Volcano Plots shows the fold change (log2 Ratio) plotted against the Absolute Confidence (-log10 adjusted p value) **(F)** Plots representing GSEA pathways in intratumoral monocytes, macrophages and DCs enriched in LysM^cre^KDM6B^fl/fl^ mice as compared to control.

To determine the impact of myeloid cell-specific *Kdm6b* deletion on the GBM tumor immune microenvironment, we performed scRNA sequencing of the murine GBM (GL261) tumors from control and LysM^cre^KDM6B^fl/fl^ mice. UMAP analyses of *Cd45+* immune cell subsets revealed distinct clusters of immune cell subsets including NK cells (C2, C18), CD8 T cells (C5,11,21), CD4 T cells (C8), regulatory T cells (C6, 22), *Il17+Rorc+* T cells (C20), B cells (C13), neutrophils (C24), mast cells (C25), microglial cells (C0 and C23), conventional dendritic cells (C3,4,7,9,17,19), plasmacytoid dendritic cells (C16) and multiple subsets of monocytic macrophages (C1,10,12,14,15) (Fig. 2C, Extended Data Fig. 7A). We noted an increase in the abundance of cytotoxic G*zmb+Ifnγ+Cd8+* T cells (C11), NK cells (C2) with a concomitant decrease in the abundance of immunosuppressive *Cd4+Foxp3+* regulatory T cells- Tregs (C6) (Extended Data Fig. 7A,B). Further, the frequency of *Il17+* T cells (C20) was lower in the TME of GBM tumors derived from LysM^cre^KDM6B^fl/fl^ mice compared to the control mice (Extended Data Fig. 7A,B). Overall, a high cytotoxic T lymphocyte (CTL) to Treg ratio in LysM^cre^KDM6B^fl/fl^ mice (Extended Data Fig.7C) indicated a pro-inflammatory skewing of the intratumoral milieu characterized by a heightened anti-tumor T cell response in LysM^cre^KDM6B^fl/fl^ mice.

The LysM-cre model used in this study harbors genetic deletion of *Kdm6b* specifically in myeloid cells ^39, 40^. Hence, the changes observed in non-myeloid immune subsets including increased T cell mediated anti-tumor immunity in the LysM^cre^KDM6B^fl/fl^ GBM (GL261) tumor-bearing mice could be secondary to the changes in *Lyz2* expressing myeloid subsets following *Kdm6b* deletion.

In our scRNA seq analysis, we noted *Lyz2* expression in intratumoral monocytic macrophages, neutrophils, certain clusters of DCs and microglial cells (Extended Data Fig.7D). To investigate the functional changes in these myeloid cell subsets, we studied the differentially expressed genes (DEGs) in these subsets from control and LysM^cre^KDM6B^fl/fl^ mice. This analysis revealed that depletion of *Kdm6b* significantly altered the transcriptomic landscape of intratumoral myeloid cells in GBM tumor-bearing mice. Expression of several pro-inflammatory genes such as those involved in phagocytosis (*Fcgr3, Lgals3, Clta, Arpc3, Arf1),* antigen presentation (*H2-Ab1, H2- Eb1*) as well as several ISGs (*Oas1a*, *Isg15, Irf7, Cxcl9* and *Cxcl10)* were upregulated whereas genes associated with immune-suppression such as *Zeb2, Klf2* and *Klf6* were downregulated in monocytic macrophages (Fig. 2D) and DCs (Fig. 2E). Next, we performed Gene Set Enrichment Analysis (GSEA) with the DEGs which revealed the differences in the major functional pathways in the intratumoral monocytes, macrophages and DCs derived from control and LysM^cre^KDM6B^fl/fl^ GBM (GL261) tumor-bearing mice. Prominent enrichment of Fc Gamma Receptor mediated phagocytosis, antigen presentation pathway as well as type I and type II interferon response was observed upon *Kdm6b* deletion (Fig.2F). In addition to the transcriptomic changes, we noted a decrease in the abundance of tumor-infiltrating neutrophils and mast cells (Extended Data Fig.7A,B). Further, we observed a concomitant increase in the abundance of pDCs (C16), a major type I interferon producing cells ^41^ and migratory cDC1s (C7) which are known to be efficient antigen-presenters ^42, 43^ in LysM^cre^KDM6B^fl/fl^ mice compared to the control (Extended Data Fig. 7A,B). Of note, migratory cDC1s express *Lyz2* while pDCs lack expression of *Lyz2* (Extended Data Fig. 7D). Hence, the changes in migratory DCs could be directly attributed to *Kdm6b* deletion, while the effect on pDCs could be due to changes in the TME. Cumulatively, the findings from scRNA seq demonstrated a global transcriptomic change in intratumoral myeloid cell subsets in the absence of *Kdm6b,* leading to a pro-inflammatory milieu in the GBM TME. Additionally, mass cytometry analysis (Extended Data Fig. 8A,B) of GBM tumors from CT-2A tumor bearing control and LysM^cre^KDM6B^fl/fl^ mice showed a decrease in the abundance of PDL1+TGFβ+ (C26) and CD115+TGFβ+ (C15) suppressive myeloid cell clusters (Extended Data Fig. 8B,C) in the absence of *Kdm6b* with a concurrent increase in the abundance of GZMB+ CD8 T cells (C22) (Extended Data Fig. 8B,C) and the CTL to Treg ratio (Extended Data Fig. 8D). Thus scRNA seq and CyTOF analysis of tumors derived from two pre-clinical GBM models demonstrated that the absence of *Kdm6b* in myeloid cells results in a pro-inflammatory milieu in murine GBM tumors.

### *Kdm6b* deletion alters the abundance and transcriptomic landscape of intratumoral monocytes and macrophages

To gain a deeper understanding of the transcriptomic changes in the highly abundant monocytes/macrophages in the GBM TME following *Kdm6b* deletion, we performed reclustering of *Itgam+* clusters C0,1,10,12,14,15,23 (Fig. 3A, Extended Data Fig. 9A). Annotation of all the distinct cell subsets demonstrated several monocytic macrophage clusters (C1-9, 11), brain-associated macrophages (C0) as well as a microglial cluster (C10) in the GBM TME (Fig. 3A, Extended Data Fig. 9A,B). Overall, we noted a decrease in the infiltration of *Chil3+Ccr2+S100a4+* classical monocytic macrophages (C7) in the LysM^cre^KDM6B^fl/fl^ tumor-bearing mice (Fig. 3B,C). Clusters expressing antigen presenting molecules (*H2-Eb1+H2-Ab1+H2-Aa+Ciita+*)(C2), interferon signature genes (*Isg15+Ifit+Irf7+Rsad2+Isg20+Tlr2+Cxcl9+Cxcl10+*)(C3) and phagocytic genes (*Lgals3+Gpnmb+Fabp5+*)(C9) were more abundant in LysM^cre^KDM6B^fl/fl^ GBM tumor-bearing mice whereas cluster expressing immunosuppressive genes (*Klf2+Klf4+Zeb2+Atf4+Mafb+Klf6+*)(C6) were more abundant in GBM tumors derived from control mice (Fig. 3B,C). In addition to cellular abundance, comparison of gene expression patterns of these five individual clusters between GBM tumors derived from control and LysM^cre^KDM6B^fl/fl^ mice showed an increase in the expression of several MHC molecules in C2 indicating enhancement of their ability to present antigens as well as increased *Myd88, Irf5, Isg20, Ifitm3, Oas1a, Isg15* in C3, indicating stronger interferon signaling in response to *Kdm6b* deletion (Fig. 3C). Further, immune-suppressive clusters C6 and C7 also showed significant upregulation of pro-inflammatory genes following *Kdm6b* deletion (Fig. 3C).

**Fig. 3:**
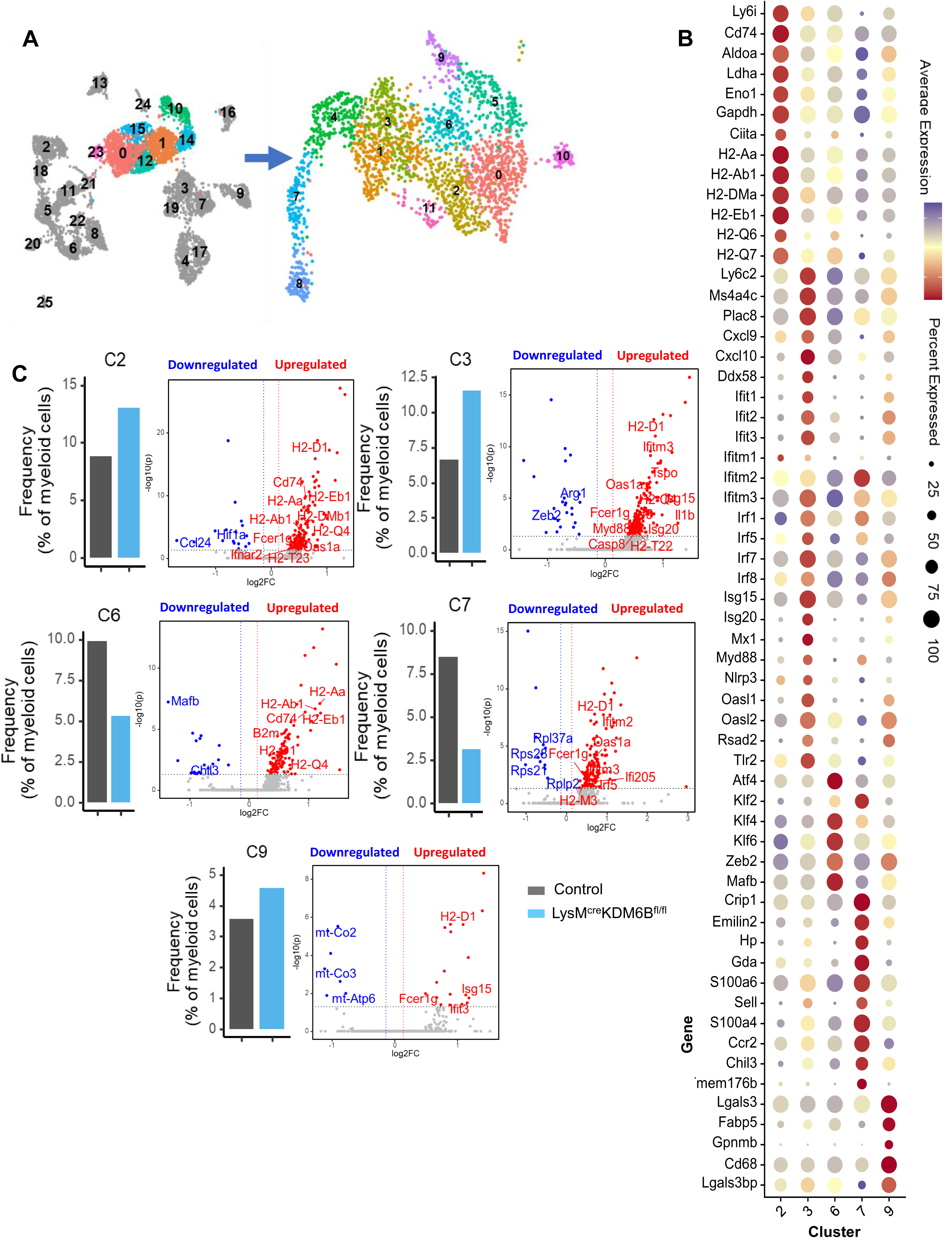
Kdm6b deletion alters the abundance and transcriptomic landscape of intratumoral monocytes and macrophages. **(A)** The left panel shows UMAP plot of scRNA seq data depicting the different immune cell subsets (*Cd45+* cells) in the GBM (GL261) TME of control and LysM^cre^KDM6B^fl/fl^ mice (pooled 4-5 samples/group, representative of two independent experiments). The right panel shows UMAP plot obtained by reclustering of highlighted *Itgam*+ clusters from the left panel. **(B)** Dotplot showing the average expression of genes of interest as well as the percentage of cells in the cluster expressing the gene in the indicated myeloid clusters. **(C)** Bar plots representing the relative frequencies of indicated myeloid clusters in control and LysM^cre^KDM6B^fl/fl^ mice. Volcano plots depicting differentially expressed genes in the indicated intratumoral myeloid cell clusters from control and LysM^cre^KDM6B^fl/fl^ mice. The Volcano Plot shows the fold change (log2 Ratio) plotted against the Absolute Confidence (-log10 adjusted p value).

Together, these findings revealed that the absence of *Kdm6b* in monocytes/macrophages leads to upregulation of pro-inflammatory gene expression, thus regulating the phenotypic plasticity of intratumoral myeloid cells.

### Myeloid cell-specific deletion of *Kdm6b* alters the chromatin landscape of key genes regulating the functional phenotype of intratumoral myeloid cells

To determine the chromatin landscape responsible for the changes in the gene expression observed in response to *Kdm6b* deletion, we performed scATAC seq of CD45*+* cells sorted from the GBM (GL261) tumors of control (n=5, pooled) and LysM^cre^KDM6B^fl/fl^ mice (n=5, pooled). Single-cell ATAC profiling of the immune cell compartment of GL261 tumors followed by UMAP analysis showed distinct immune cell subsets (Fig. 4A,B, Fig. S10A). Genes proximal to cluster-specific cis-elements were also used to annotate individual cell types (Fig. 4C). Briefly, C(0-2,6) showed accessibility at cis-elements neighboring macrophage associated genes, including *Mafb, Cebpb* and *F10*; C(10,13) demonstrated accessible cis-elements proximal to DC associated genes such as *Mreg, Nr4a3* and *Anxa3*, while C(5,14) harbored accessible cis-elements neighboring B cell associated genes, including *Fam43a, Cd19* and *Ms4a1* (Fig. 4C). Additionally CD4 T cells (*Cd4, Tcf7, Zap70*), CD8 T cells (*Cd8a, Cd8b, Ifng*), regulatory T cells (*FoxP3*), NK cells (*Eomes, Prf1*) and even rare cell subsets such as mast cells (*Homer2, Tbc1d8*) could be identified from accessibility profiles of cell type specific cis-regulatory elements (Fig. 4C). We also noted a prominent increase in the abundance of CTLs (C12) and a concomitant decrease in the abundance of Tregs (C21) (Extended Data Fig.10 A,B), mirroring the findings from scRNA seq and confirming the pro-inflammatory skewing of the intratumoral milieu in LysM^cre^KDM6B^fl/fl^ mice.

**Fig. 4:**
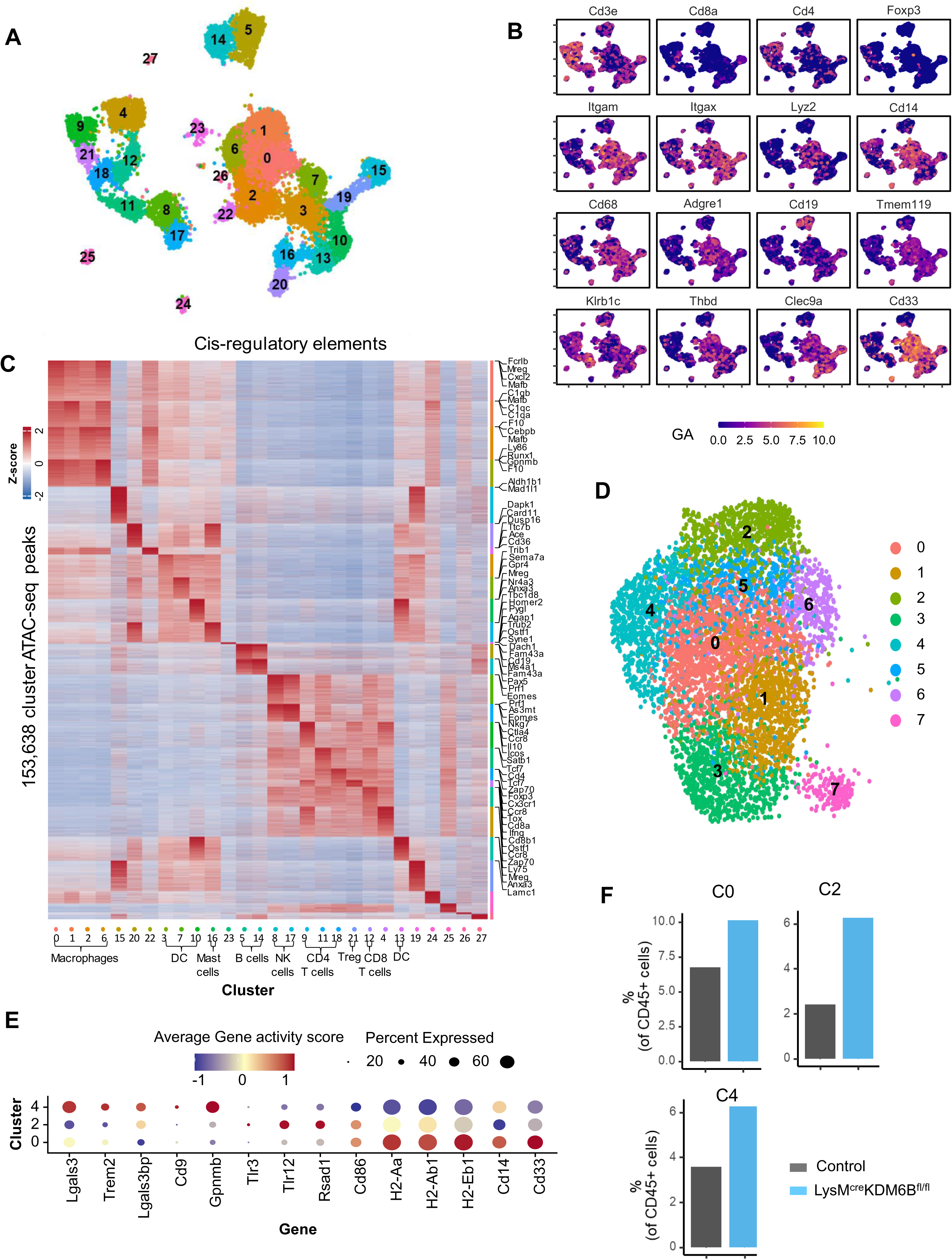
Myeloid cell-specific deletion of Kdm6b alters the chromatin landscape of key genes regulating the functional phenotype of intratumoral myeloid cells. **(A)** UMAP demonstrating the different Cd45+ immune cell subsets in the GBM (GL261) TME of control and LysM^cre^KDM6B^fl/fl^ mice as determined from scATAC seq (4-5 pooled samples/group). **(B)** UMAP depicting the gene activity score of the indicated genes in different immune cell subsets in the GBM TME of control and LysM^cre^KDM6B^fl/fl^ mice. **(C)** Heatmap showing Z-scores of 153,638 cis-elements neighboring indicated genes in the scATAC seq clusters derived from Fig.4A. **(D)** UMAP representation of scATAC seq data depicting Itgam+ cell subsets in the GL261 TME of control and LysM^cre^KDM6B^fl/fl^ mice (pooled 4-5 samples/group). **(E)** Dotplot demonstrating the average gene activity score of genes of interest as well as percentage of cells in the cluster expressing the gene in the indicated myeloid cell clusters. **(F)** Bar graphs representing the relative frequencies of indicated myeloid clusters from control and LysM^cre^KDM6B^fl/fl^ mice.

To define the chromatin landscape of myeloid cells, we analyzed *Lyz2+* population which showed an increase in abundance of antigen presenting cluster (C0), cluster expressing IFN-related genes (C2) as well as the phagocytic cluster (C4) (Fig. 4D-F, Extended Data Fig.11A). In addition to the quantitative changes observed in the myeloid cell populations, we aimed to address the qualitative changes occurring in the different myeloid subsets in response to *Kdm6b* deletion. Interrogation of chromatin accessibility of individual genes of interest via coverage plots revealed greater accessibility of genes associated with antigen presentation such as *H2-Eb2, H2-Ab1* (C0), genes encoding positive regulators of phagocytosis such as *Fcgr1* and *Fcgr4* (C0) (Extended Data Fig.11B), and genes involved in interferon signaling and response such as *Ifnar1, Ifngr1, Isg15, Ifitm6* (C2) in LysM^cre^KDM6B^fl/fl^ mice (Extended Data Fig.11C).

Thus, the findings from scATAC seq showed open chromatin landscape of pro-inflammatory genes in intratumoral myeloid cells in the absence of *Kdm6b*.

### *Kdm6b* regulates H3K27me3 enrichment of genes regulating phagocytosis, antigen presentation, and interferon response in myeloid cells

As we previously mentioned, KDM6B promotes gene expression by demethylation of H3K27me3^33^. Therefore, to test whether the observed changes in the chromatin accessibility and gene expression leading to pro-inflammatory skewing of myeloid cells were regulated by KDM6B- mediated H3K27me3 demethylation, we performed ChIP seq assays on bone marrow derived macrophages (BMDMs) from control and LysM^cre^KDM6B^fl/fl^ mice. ChIP using anti-KDM6B antibody allowed identification of genes directly bound by KDM6B. We identified *Socs3* and *Mafb* as direct targets of KDM6B in control BMDMs (Fig. 5A,D). Further, in BMDMs harbouring *Kdm6b* deletion there was a drastic reduction in KDM6B occupancy of these genes as expected (Fig. 5A,D), and a concurrent enrichment of H3K27me3 (Fig. 5B,E). We also used quantitative PCR to confirm the reduction in expression of these genes in LysM^cre^KDM6B^fl/fl^ BMDMs (Fig. 5C,F). These findings demonstrated that KDM6B directly binds to *Socs3* and *Mafb* encoding genetic regions and demethylates H3K27me3 to induce gene expression. Hence in the absence of KDM6B, these genes are enriched for the repressive H3K27me3 marks thus inhibiting gene expression. SOCS3 is a known suppressor of cytokine signalling^34^ and MAFB has been established as a suppressor of type I IFN signalling^35^. Therefore, reduced expression of these immune-suppressive genes in the absence of KDM6B provides a strong rationale for the pro-inflammatory skewing of myeloid cells observed in response to *Kdm6b* deletion. Further, ChIP seq identified *Sirpa* as a direct target of KDM6B, with reduction in KDM6B occupancy (Fig. 5G) and H3K27me3 enrichment (Fig. 5H) in *Kdm6b* deleted BMDMs. SIRPA acts as a negative regulator of phagocytosis by generating “don’t-eat-me” signals^19^. Overall, the findings from the ChIP-sequencing study provided mechanistic insight into KDM6B mediated regulation of macrophage phenotype and function. KDM6B binds to negative regulators of interferon response and phagocytosis including *Mafb, Socs3* and *Sirpa*. Thus, these findings implicate KDM6B as an upstream regulator of multiple critical functional pathways in macrophages including cytokine production/response, antigen presentation and phagocytosis.

**Fig. 5:**
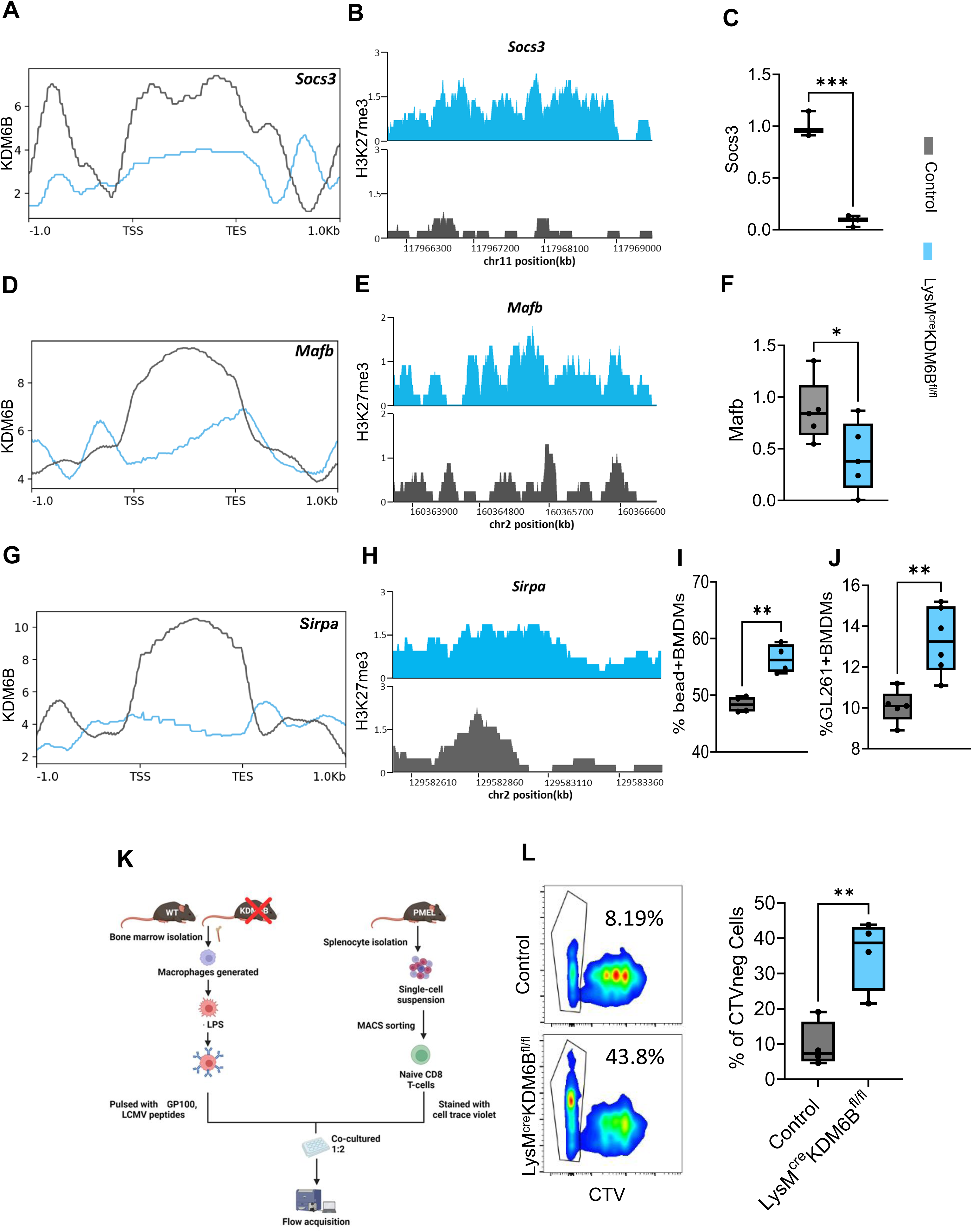
Kdm6b regulates H3K27me3 enrichment of genes regulating phagocytosis, antigen presentation, and interferon response in myeloid cells. **(A, D, G)** Profile plots depicting the probability scores of KDM6B binding at -/+1kb regions from transcription start site (TSS) and transcription end site (TES) of the Socs3, Mafb and Sirpa gene loci in bone marrow derived macrophages (BMDMs) from control (grey) and LysM^cre^KDM6B^fl/fl^ (blue) mice. **(B, E, H)** Genome browser view of H3K27me3 peaks at the Socs3, Mafb and Sirpa gene loci in control and LysM^cre^KDM6B^fl/fl^ BMDMs. **(C, F)** Box and whisker plots showing the relative expression of indicated genes normalized to the expression of β-actin as determined by quantitative PCR. One- tailed Student’s t-test was performed (*p<0.05, ***p<0.001, n=3-6/condition). **(I,J)** Box and whisker plots representing the difference in phagocytic capacity between control and Kdm6b deficient BMDMs, based on percentage of cells taking up beads **(I)** or GL261 cells **(J)**. Two-tailed Student’s t-test was performed (**p<0.01). n=5-6/group, representative of 2-4 independent experiments. **(K)** Schematic representation of the antigen presentation and T cell proliferation assay performed. **(L)** Representative pseudocolor flow cytometry plots showing the percentage of proliferated CD8 T-cells (gated, CTV negative), upon co-culture with gp100 pulsed control and Kdm6b deficient BMDMs. Box and whisker plot depicting the percentage of proliferated CD8 T- cells upon co-culture with control versus Kdm6b deficient macrophages. (n=4, **p<0.01). Data is representative of two independent experiments.

In order to investigate the impact of KDM6B depletion on phagocytosis and antigen-presentation functions of myeloid cells, we performed in-vitro phagocytosis and antigen-presentation assays using bone marrow derived macrophages (BMDMs) from control and LysM^cre^KDM6B^fl/fl^ mice. We found that following stimulation with lipopolysaccharide, *Kdm6b* deficient BMDMs demonstrated enhanced phagocytosis compared to control as evident from a higher percentage of fluorescent non-coated latex bead positive BMDM cells (Fig. 5I). Additionally, phagocytosis of fluorescently labelled GL261 cells was higher by BMDMs deficient in *Kdm6b* compared to control (Fig. 5J, Extended Data Fig.11D). For antigen presentation assay we co-cultured gp100-pulsed BMDMs with cell trace violet (CTV) labelled cognate T cell receptor-bearing CD8 T cells isolated from pmel mice^44^ (Fig. 5K). Based on differences in dilution of the CTV dye, we observed that co-culture with BMDMs derived from LysM^cre^KDM6B^fl/fl^ mice led to significantly higher T cell proliferation as compared to BMDMs derived from control mice (Fig. 5L). BMDMs pulsed with the non-cognate LCMV peptide, showed minimal proliferation thus confirming the antigen-specificity of the observed proliferation in T cells (Extended Data Fig.11E). Overall, the T cell proliferation assay indicated that *Kdm6b* deficient BMDMs are more efficient antigen presenters as compared to control BMDMs. These findings provided evidence of enhanced phagocytic and antigen presentation function of myeloid cells following *Kdm6b* deletion.

Cumulatively, we found that KDM6B regulates H3K27me3 enrichment of genes regulating critical pathways modulating myeloid cell functions such as phagocytosis, antigen presentation, and interferon signaling/response.

### Pharmacological inhibition of KDM6B improves the efficacy of immune checkpoint therapy in GBM

Since KDM6B functions upstream to several critical functional pathways, therapeutic targeting of KDM6B could revert myeloid-derived immune suppression. To determine the translational relevance of our findings from the genetic model, we compared murine GBM tumor growth and the tumor immune microenvironment in the presence and absence of a pharmacological inhibitor of KDM6B (GSK-J4)^45, 46^. MRI studies revealed lower tumor burden in GSK-J4 treated GL261 tumor bearing mice as compared with control (Fig. 6A). Additionally, we observed improvement in the overall survival of GL261 tumor-bearing mice treated with GSK-J4 (Fig. 6B). Of note, GSK- J4 which targets KDM6A/B, has been shown to inhibit proliferation of glioma cell lines in-vitro^47^.

**Fig. 6:**
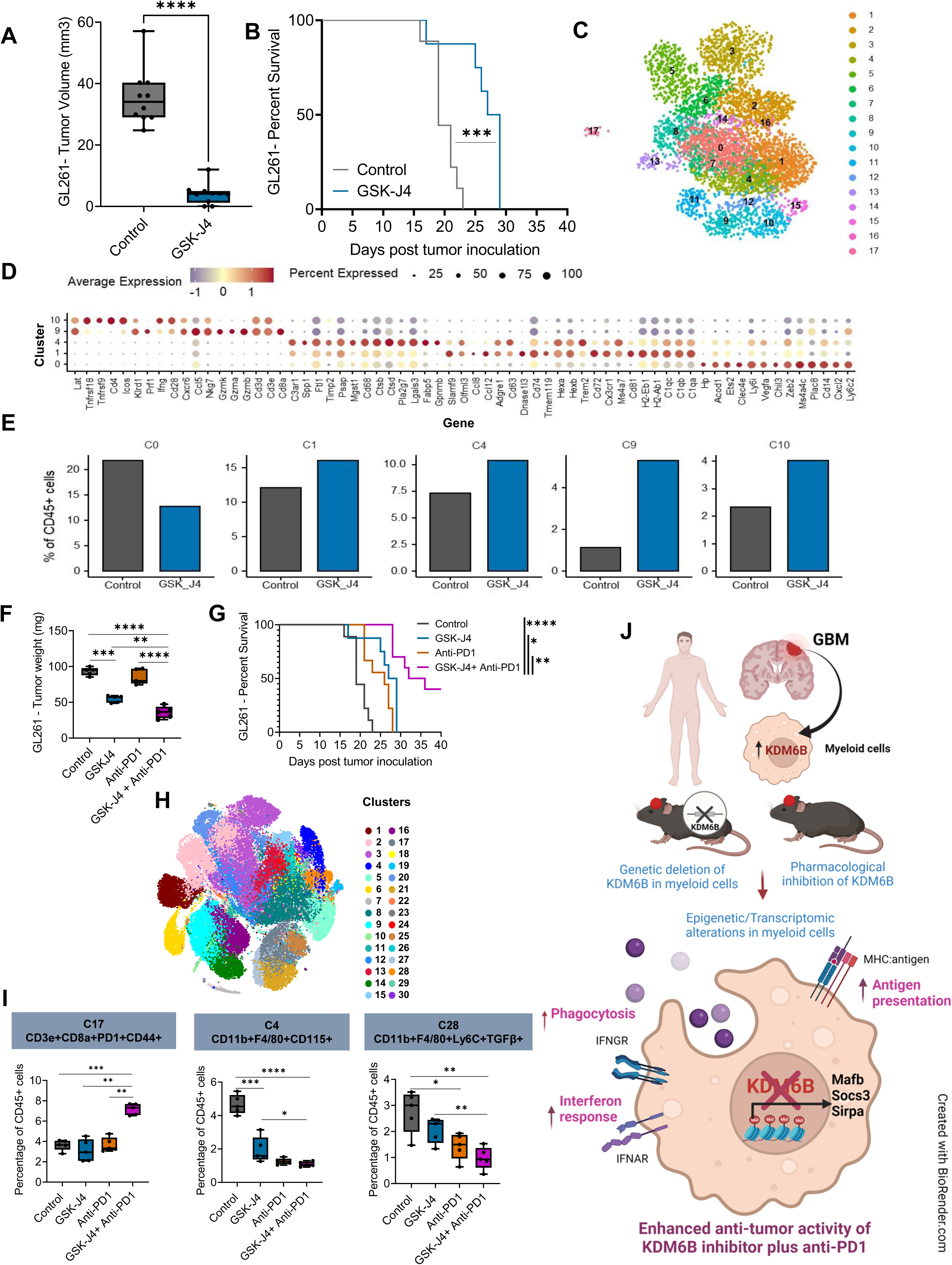
Pharmacological inhibition of KDM6B improves the efficacy of immune checkpointtherapy in GBM. **(A)** Box and whisker plot depicting the difference in the tumor volume as calculated from day 14 MRI images of GBM (GL261) tumors from control (vehicle treated) and GSK-J4 treated mice (n=10/group). Two-tailed Student’s t-test was performed (**** p<0.0001). **(B)** Kaplan Meier plot demonstrating the difference in survival of GBM (GL261) tumor-bearing mice treated with vehicle and GSK-J4 (n=10/group). Log-rank test was performed (*** p<0.001). Data is representative of 3 independent experiments. **(C)** UMAP plot of scRNA seq data representing the different immune cell subsets (*Cd45*+ cells) in the GBM (GL261) TME of vehicle and GSK-J4 treated mice (pooled 4-5 samples/group). **(D)** Dotplot showing the average expression of genes of interest as well as percentage of cells in the cluster expressing the genes defining the indicated cell clusters. **(E)** Bar plots representing the frequencies of indicated immune cell clusters from vehicle and GSK-J4 treated mice as determined by scRNA seq. **(F)** Box and whisker plot representing the GBM (GL261) tumor weights from mice receiving the indicated treatments (n=5/group). Two-tailed Student’s t-test was performed (**p<0.01, ***p<0.001, ****p<0.0001). **(G)** Kaplan Meier plot depicting difference in survival of GBM (GL261) tumor-bearing mice treating with vehicle, anti-PD1, GSK-J4 and combination of anti-PD-1 plus GSK-J4. Log-rank test was performed (*p<0.05, **p<0.01, ****p<0.0001). Data is representative of 2 independent experiments. **(H)** t-SNE plot of CyTOF data demonstrating different immune cell subsets (CD45+ cells) in the GBM (GL261) TME of vehicle, GSK-J4 and anti-PD1 treated mice (n=5/group). **(I)** Box and whisker plots showing the relative frequencies of indicated immune cell clusters from vehicle treated and therapeutic agent treated mice as determined by CyTOF (n=5/group). Two- tailed Student’s t-test was performed (*p<0.05, **p<0.01, ***p<0.001,****p<0.0001). **(J)** Graphical summary of the findings presented in this study depicting the role of KDM6B in regulation of myeloid cell functionand its importance as a therapeutic target in GBM. IFNGR- Interferon gamma receptor. IFNAR- Interferon alpha receptor. GBM- Glioblastoma.

Hence, the observed improvement in survival could be due to the direct effect of GSK-J4 on tumor cells as well as its effects on the tumor immune microenvironment.

Hence, to investigate the effect of GSK-J4 on the GBM tumor immune microenvironment, we performed scRNA seq. UMAP analyses performed on scRNA-seq data from GSK-J4 treated and control GL261 tumors showed distinct lymphoid (C9-12,15,17) and myeloid cell clusters (C0- 8,13,14,16) (Fig. 6C,D, Extended Data Fig. 12). We noted a significant decrease in the abundance of *Cd14+Ly6c2+Plac8+Cxcl2+Vegfa+* monocytic macrophage subset (C0) in GBM tumor-bearing mice treated with GSK-J4 (Fig. 6E). C1, characterized by the expression of markers of both activated CNS associated macrophages and disease associated microglia (*Ms4a7, Ccl8, Cd74, Tmem119, Hexb, Cx3cr1, Trem2, C1qa, C1qb, C1qc, H2-Ab1*) increased in abundance post treatment with GSK-J4 (Fig. 6D,E). We also identified a *Gpnmb+* phagocytic cluster (C4) which increased in abundance in GSK-J4 treated mice (Fig. 6D,E). Importantly, we also noted an increase in the *Cd8+Gzmb+Ifng+* cytotoxic T cells (C9) and *Cd4+Icos+Ifng+Cxcr6+* T cell cluster (C10) in response to GSK-J4 treatment (Fig. 6D,E).

To corroborate the findings from the GL261 model, we used the CT-2A GBM model and treated them with GSK-J4. MRI studies revealed lower tumor burden in GSK-J4 treated CT-2A tumor bearing mice as compared with control (Extended Data Fig.13A-C). Additionally, interrogation of the TME of CT-2A GBM tumors using CyTOF (Extended Data Fig.13D,E) also showed significantly lower abundance of immune-suppressive myeloid clusters (C3,9) and higher abundance of NK cells (C12) and effector/memory CD8 T cells (C6) in GSK-J4 treated mice (Extended Data Fig.13F). Thus cumulatively, scRNA seq and CyTOF data from two distinct GBM tumor bearing murine models revealed a significant reduction in intratumoral immune-suppressive monocytic macrophage populations with a concomitant increase in effector CD8 T cell subset in response to GSK-J4 treatment.

Myeloid heavy tumor types such as GBM tumors often demonstrate primary resistance to immune checkpoint therapy. To test if KDM6B inhibition mediated pro-inflammatory skewing of the tumor immune microenvironment could increase the efficacy of ICT in a murine model of GBM, we treated GL261 tumor-bearing mice with vehicle, anti-PD1, GSK-J4 and the combination of anti- PD1 plus GSK-J4. We found that the combination therapy of anti-PD1 plus GSK-J4 led to a significant reduction in tumor weight (Fig. 6F) as well as in improvement in overall survival (Fig. 6G). In order to understand the changes in the TME, we performed CyTOF analysis on these tumors (Fig. 6H, Extended Data Fig.14). We noted that CD8+CD86+CD44+ effector memory T cell cluster (C17) was significantly higher in mice receiving a combination of anti-PD1 plus GSK- J4 as compared with the control, GSK-J4 and anti-PD1 monotherapy groups (Fig. 6I, Extended Data Fig.14). Also, there was a significant decrease in a CD11b+F4/80+ monocytic macrophage cluster expressing CD115/CSF1R (C4) and a Ly6c+Ly6g-CD11b+F4/80+ monocytic-myeloid derived suppressor cell (M-MDSCs) cluster expressing TGFβ (C28) following treatment with the combination of anti-PD1 plus GSK-J4 (Fig. 6I, Extended Data Fig.14).

Together, these findings demonstrate that pharmacological inhibition of KDM6B by GSK-J4 can skew the TME of GBM tumor-bearing mice to a pro-inflammatory phenotype and reduce the frequency of several pro-tumorigenic myeloid cell subsets including M-MDSCs and *Cxcl2+* TAMs, thus, improving overall survival and enhancing sensitivity to anti-PD1 therapy.

## Discussion

This study identified a selective expression of KDM6B in the intra-tumoral myeloid cell subsets in the GBM tumors resected from patients utilizing single-cell transcriptomic and spatial analysis. Reverse translational studies using pre-clinical models of GBM demonstrated that targeting KDM6B-mediated epigenetic pathways in the myeloid cells via genetic deletion and pharmacological inhibition resulted in upregulation of pro-inflammatory pathways, cumulatively improving survival and enhancing sensitivity to anti-PD1 therapy (Fig. 6J).

Over the years, multiple myeloid cell-specific pathways have been explored individually as potential targets for the treatment of cancer ^15–17^. However, these pathways are highly interrelated and often redundant, therefore targeting a single pathway often fails to elicit optimum clinical benefit. We found that KDM6B is upstream of multiple pathways and inhibition of KDM6B enhanced interferon response pathways, antigen presentation, and phagocytosis in macrophages as well as in DCs. Additionally, our mechanistic studies identified certain critical regulators of the above mentioned pathways including *Mafb, Socs3* and *Sirpa* as direct targets of KDM6B mediated H3K27me3 demethylation. MAF BZIP Transcription Factor B (MAFB) which encodes a basic leucine zipper myeloid cell-specific transcription factor acts as a rheostat to inhibit type I IFN induction by physically blocking IRF3 from binding to its target genes ^35, 48, 49^, which might provide a possible explanation for the enhanced interferon response observed upon KDM6B deletion. Further, suppressor of cytokine signaling 3 (Socs3) inhibits cytokine-induced JAK-STAT signaling pathways^50^ and cytokines including IFN pathways have been shown to regulate both antigen presentation and phagocytosis by myeloid cells ^51–55^. Additionally, KDM6B regulates *Sirpa* which acts as a negative regulator of phagocytosis by generating “don’t-eat-me” signals ^56^. Thus, KDM6B regulates phagocytosis by modulating both activators and inhibitors of the phagocytosis pathway. Of note, the changes observed in the phagocytosis assays, though statistically significant were quantitatively modest. This might possibly be due to the in-vitro setting in which the experiments were performed, as opposed to the in-vivo intratumoral environment. Overall, our findings suggest that KDM6B functions upstream of several critical functional pathways (Fig. 6J). Therefore, targeting KDM6B to reprogram the immune-suppressive myeloid population into an immune-stimulatory phenotype could potentially be an important therapeutic strategy rather than targeting individual pathways.

Considering the heterogeneity of the myeloid cell subsets, a single cre-flox model system can not target all the myeloid cell subsets simultaneously. We used the LysM-cre model to evaluate KDM6B-mediated epigenetic regulation of myeloid cells since *Lyz2* expression is generally observed in monocytes and macrophages, granulocytes and in some dendritic cells ^57^. Although we noted an increase in the microglial-like cells expressing antigen presentation molecules following deletion of KDM6B, LysM-cre model is not adequate for a comprehensive interrogation of microglial cells.

Overall, based on the single-cell and spatial transcriptomic analysis of human GBM samples and the series of reverse translational studies using multiple pre-clinical model systems, we identified a KDM6B-mediated immunoregulatory program in myeloid cells, providing a strong rationale to consider evaluating KDM6B inhibition as a therapeutic strategy to overcome myeloid-derived immune suppression and enhance response to immune-based therapies. The strategy of inhibiting KDM6B, proposed in this study, not only adds to the existing repertoire of myeloid cell targeting strategies, it proposes a new paradigm of regulating the epigenetic machinery to target intratumoral myeloid cell plasticity thus reprogramming them to a pro-inflammatory phenotype.

## Methods

### Patients

Patient samples were collected after appropriate informed consent was obtained on MD Anderson internal review board (IRB)-approved protocol PA13-0291. All patients signed informed consents for participation in PA13-0291 before surgery or sample collection. The clinical characteristics of individual patients are shown in Supplementary Table 1.

### H&E and IHC staining

Hematoxylin and Eosin (H&E) and IHC staining were performed on formalin-fixed, paraffin-embedded tissue sections. Tissues were fixed in 10% formalin, embedded in paraffin, and sectioned at four-micron–thickness. For IHC, sections were antigen retrieved with ER solution 1 (Leica Microsystems, catalog no. AR9961), protein block was applied for 30 mins (Leica Microsystems, catalog no. RE7102) and stained with KDM6B (ThermoFisher, catalog no. PA5-32192) at 1:200 dilution followed by rabbit anti-human secondary. 3′-3-diaminobenzidine (DAB) substrate (Leica Microsystems) was used as a chromogen followed by hematoxylin counterstain. Slides were scanned and digitalized using the scanscope system from Scanscope XT, Aperio/Leica Technologies. IHC staining was interpreted in conjunction with H&E stained sections.

### Immunofluroscence

Using the Opal multiplex immunofluorescence staining protocol ^54^ on a RX- BOND (Leica) autostainer, GBM tissue sections were stained for CD3 (Dako, A0452, 1:200 dilution), CD8 (LS-Bio, C8/144b, 1:100 dilution), CD68 (Dako, PGM-1, 1:25 dilution) and CD163 (Leica Microsystems, 10D6, 1:20 dilution). Subsequent visualization was performed using Akoya Opal fluorophores (620, 520, 480, 570 respectively), DAPI (1:2000 dilution) and cover-slipped using Vectashield Hardset895 mounting medium. Slides were scanned using a Vectra/Polaris slide scanner (PerkinElmer) and images acquired at 20X magnification were spectrally unmixed using Inform software (Akoya).

### Spatial Transcriptomics Assay (10X Genomics, Visium)

GBM tumors were paraffin embedded and serially sectioned (thickness 5 μm). Formalin-fixed paraffin-embedded (FFPE) tissue from 3 GBM samples were used for spatial transcriptomics analysis. FFPE samples were tested for RNA quality with an DV200 > 30% (Agilent). The samples were then processed according to the standard Visium Spatial Gene Expression protocol (10x Genomics) using the Visium Spatial Gene Expression Slide & Reagent Kit (10x Genomics).

Libraries were cleaned up using SPRI select reagent and quantified using the High Sensitivity DNA Kit run on Agilent 2100 Bioanalyzer and also KAPA Illumina library quantification kit (Roche, 7960140001) run on LightCycler 480. Library pool was quantified on Bioanalyzer and with quantitative PCR and sequenced using Illumina NextSeq 500.

### Visium spatial transcriptomics data analysis using Spaceranger and CellTrek

The raw spatial sequencing data was processed in the Spaceranger workflow (10X Genomics). The spaceranger (version 2.0.0) mkfastq pipeline was used to convert Illumina sequencer’s binary base call (BCL) files into FASTQ format. Samples were then run through the spaceranger count pipeline, which performs alignment, tissue detection, fiducial detection, and barcode/unique molecular identifier counting. Human GBM scRNA seq data analyses were performed using standard Seurat data analysis pipeline including log normalization, scaling, variable genes selection (n=3,000) using vst, dimension reduction using principal component analysis (PCA) and UMAP. Spatial transcriptomics (ST) data (10x Genomics Visium) was analyzed similar to the scRNA seq data with Seurat data analysis package (log normalization, scaling, variable genes identifcation and dimensionality reduction). To analyze ST data using CellTrek ^38^, we first ran traint to coembed the data into a shared feature space with default parameters. Then we ran Celltrek on the coembedded traint data with following parameters: intp_pnt =5,000 spots, nPCs =30, ntree =1,000, top_spot =5, spot_n =5 and repel_r =3 with ten iterations. To visualize the ST data, we used celltrek_vis tool which allows mapping any continuous or categorical cell features to the spatial map with different colors.

### Mice

5–7-weeks old C57BL/6 mice were purchased from the National Cancer Institute (NCI). 5– 7-weeks old KDM6b^fl/fl^(B6.Cg-Kdm6b^tm1.1Rbo^/J, Stock no. 029615) mice and LysM^cre^ (B6.129P2- Lyz2^tm1(cre)Ifo^/J, Stock no. 004781) mice were purchased from the Jackson Laboratory. Female mice were used for the experiments. All mice were kept in specific pathogen-free conditions at the Animal Resource Center, University of Texas MD Anderson Cancer Center. Animal protocols were approved by the Institutional Animal Care and Use Committee of the University of Texas MD Anderson Cancer Center.

LysM^cre^Kdm6b^fl/fl^ mice were generated by breeding KDM6b^fl/fl^ and LysM^cre^ mice ^39, 40^. PCR based genotyping study was done to confirm *Kdm6b* deficiency using primers with the following sequences Forward- 5’-CAG CGA TCC TGA CTT GTT CA-3’ Reverse- 5’-GTG CCA AGG CTG GAG GA-3’

### Mass cytometry based immunophenotyping assay

Spleen, bone marrow and lymph node were collected from control and LysM^cre^Kdm6b^fl/fl^ mice. Single-cell suspensions were generated by physical dissociation and passage through 70μm filters. Cells were washed in RPMI media by centrifugation at 2,000 rpm, 4 °C for 5 minutes. Upto 3× 10^6^ cells were washed with FACS buffer containing 5% FBS (Fetal Bovine Serum) in PBS (Phosphate Buffered Saline) and then incubated with blocking buffer containing a mixture of 2% of bovine, murine, rat, hamster, rabbit serum and 25 μg/mL 2.4G2 antibody (Fc block) in PBS at 4 °C for 10 min. Next, surface staining was done with antibody mixture (Supplementary Table 2) at 4 °C for 30 min. Following incubation,194Pt monoisotopic cisplatin (Fluidigm) in PBS at a final concentration of 5 μM, was incubated with the samples for 3 min. Next, samples were washed twice with FACS buffer followed by fixation and permeabilization for 1 hour at 4°C. After a wash with permeabilization buffer (Invitrogen) intracellular staining was done for 30 mins at 4°C (Supplementary Table 2). Following staining, samples were washed twice with Maxpar barcode perm buffer (Fluidigm) and labeled using palladium barcoding as per the manufacturer’s protocol for 30 min at room temperature. Following 2 washes with FACS buffer, samples were fixed using 1.6% paraf ormaldehyde in PBS supplemented with 100 nM iridium nucleic acid intercalator (Fluidigm) and left overnight. Next day, cells were washed twice with PBS, filtered, and resuspended in nuclease free water. Barcoded samples were then acquired in a Helios mass cytometer (Fluidigm).

### Cell lines and tumor models

The murine GBM cancer cell line GL261 was obtained from NCI and the CT-2A cell line was obtained from Millipore Sigma. GL261 and CT-2A cells were cultured in complete (supplemented with 10% FBS) DMEM media at 37 °C and 5% CO_2_. Cells in the logarithmic phase of growth were harvested by trypsinization and washed twice with PBS before intracranial inoculation. 5X10^4^ GL261 cells were resuspended in 3µl of 70% PBS and 30% Matrigel while 5X10^4^ CT-2A cells were resuspended in 3µl of DMEM media (without FBS) for injection with a stereotactic apparatus (Stoelting) in the cranial window- 2mm posterior and 2mm lateral to the bregma and 3mm deep into the mouse cerebrum.

### MRI image quantification

The MRI images were quantified using ImageJ (NIH) v.1.52a. First, images were imported, and their brightness/contrast was adjusted. Image slices were then scanned to identify tumor sections. A gate was drawn around the tumor in each section and the area was measured. Image geometry indicated the slice thickness to be 0.75 mm and the distance between two sections to be 1 mm. The tumor area in each section was multiplied by 0.75 and the average between the tumor area in two sections was taken and multiplied by 0.25 (1 − 0.75; this gave the value for depth). The volume for each tumor was obtained by multiplying the tumor area and depth from the section-containing tumor. All values were added to determine tumor volume in mm^3^.

### Tumor harvesting and processing

GL261 tumors were harvested on day 17 post tumor inoculation and CT-2A tumors were dissected on day 22 post tumor inoculation. Following dissection, tumor samples were enzymatically digested with 0.66mg/ml Liberase TL (Roche) and 20mg/ml DNase I (Roche) in RPMI cell culture media for 30 minutes at 37 °C. Single-cell suspensions were generated by passing digested tumors through 70-μ filters and washed in complete RPMI media by centrifugation at 2,000 rpm 4°C for 5 minutes. Percoll gradient centrifugation at 512g for 20 minutes at 18°C was used to deplete the myelin layer and the single cell suspension obtained was counted in an automated cell counter for downstream analysis.

### Single cell RNA sequencing

Single cell suspension of human and murine GL261 GBM tumors were made using the protocol described above. Single cells were incubated with a surface staining cocktail of fluorescently conjugated antibodies, which included CD45 Pacific Blue (clone 30-F11, Biolegend, 103126), and live/dead discrimination viability dye Pacific Orange (Invitrogen, L34968). CD45+ cells were sorted directly into 5% FBS using a FACS AriaFusion cell sorter (BD). Cells from each sample were counted before 16,000 cells per sample were loaded on the 10x chromium chip (Chromium platform,10x Genomics), with a target of 10,000 cells per sample for the downstream analysis. Single-cell mRNA libraries were built using the Chromium Next GEM Single Cell 3’ Library Construction V3 Kit, libraries sequenced on NovaSeq 6000 using 100cycle kit, flow cell type – S2-100, run format- 28/91 and 8 i7 index.

### Single cell RNA sequencing data analysis

Cellranger v3.0.2 software (10x Genomics) was used to process the sequencing reads. The “cellranger count” pipeline was used to align the reads to the mouse mm10 genome and compute the count matrix. The Seurat R package was used to perform the analysis including filtering out low-quality cells, normalizing the data, and clustering the cells. Genes presented in less than 3 cells and cells with less than 200 genes or more than 6000 genes, or with more than 10% mitochondrial gene counts were excluded from downstream analysis. Potential doublets were removed with the DoubleFinder R package ^58^. Then, the “SCTransform” function was used to normalize and log transform the raw gene counts. Anchors identified by the “FindIntegrationAnchors” function were used to integrate the datasets. Principal component analysis (PCA) was applied to the top 3000 highly variable genes and the first 30 components were used for constructing a KNN graph, clustering and UMAP projection.

### Single cell ATAC sequencing

GL261 tumors were dissociated from control, LysM^cre^Kdm6b^fl/fl^ mice and single cell suspension of cells were made using the protocol described above. Single cells were incubated with a surface staining cocktail of fluorescently conjugated antibodies, which included CD45 Pacific Blue (clone 30-F11, Biolegend, 103126), CD3ε FITC (clone 17A2, eBioscience, 11-0032-82), CD11b APC (clone M1/70, Biolegend, 101212) and live/dead discrimination viability dye Pacific Orange (Invitrogen, L34968). CD45+ cells were sorted directly into 5% FBS using a FACS AriaFusion cell sorter (BD). Cell nuclei were isolated from the sorted CD45+ cells using Nuclei Isolation for Single Cell ATAC Sequencing Protocol (CG000169 Rev D). 16,000 nuclei per sample were loaded on the 10x chromium chip (Chromium platform,10x Genomics), with a target of 10,000 nuclei per sample for the downstream analysis. Single cell ATAC libraries were built according to the manufacturer’s protocol (Chromium Next GEM Single Cell ATAC Reagent Kits (v1) User Guide - CG000168 Rev D). The libraries were pooled and sequenced using NovaSeq6000 instrument with Read1 (50 cycles), Read2 (49 cycles), Index1 (8 cycles), and Index2 (16 cycles). The sequencing reads were demultiplexed based on sample index barcodes.

### Single cell ATAC sequencing data analysis

Cellranger-atac v1.2 software (10x Genomics) was used to process the sequencing reads. The “cellranger-atac count” pipeline was used to align the reads to the mouse mm10 genome and to perform peak calling with the default parameters. In case the detected cell number from the auto cell detecting algorism was unexpected, the “force-cells” parameter for the “cellranger-atac count” was manually set according to the Barcode Rank Plot in the web summary result. Peak barcode count matrices from all the samples were aggregated using the “cellranger-atac aggr” pipeline function and normalized to sequencing depth. The single-cell ATAC data analysis mainly was done using the Signac (version 1.1.0) (https://github.com/timoast/signac) and Seurat ^59^(version 3.2.0) R packages.

Cluster-wise peak calling was performed with MACS2 as previously described^60^. Briefly, the mouse genome was tiled into 2.5 kb size windows and a cell-by-window sparse matrix was computed by the Signac “FeatureMatrix” function. The matrix was binarized and the top 20000 most accessible sites across all cells were used to cluster the cells. Peak calling for each cluster was performed by the Signac “CallPeaks” function and a union peak set of 180413 was created. Then, a cell-by-peak sparse count matrix was computed by the Signac “FeatureMatrix” function and used for downstream analysis.

Peaks presented in less than 10 cells and cells with less than 200 peaks were removed from the downstream analysis. Quality control matrixes including percentage reads in peaks, blacklist ratio, nucleosome signal and TSS enrichment score were calculated following the Signac vignettes. Cells with peak region fragment count between 3000 ∼ 50000, percentage reads in peaks > 50, blacklist ratio < 0.025, nucleosome signal < 4, and TSS enrichment score > 2 were considered for further analysis (total 17380 cells).

The peak barcode count matrix was binarized and was normalized by term frequency-inverse document frequency (TF-IDF) using the Signac “RunTFIDF” function. Dimensional reduction was performed with a singular value decomposition (SVD) on the TD-IDF normalized matrix using the Signac “RunSVD” function including all the peaks. Since the first LSI component usually captures the sequencing depth variation, it was removed for the downstream analysis. Graph-based clustering, and non-linear dimension reduction for visualization were performed using the Seurat “RunUMAP”, “FindNeighbors”, and “FindClusters” functions with the 2 to 50 LSI components and resolution of 1.4. To find differentially accessible perks between two groups of cells, the Seurat “FindMarkers” function was used to with the parameter “test.use = “LR”, latent.vars = “peak_region_fragments”. The identified peaks was annotated on the basis of its nearest gene using the Signac “ClosestFeature”. Peak visualization was performed with the Signac “CoveragePlot” function.

Gene activity score was calculated using the Cicero (version 1.3.4) and Monocle3 (version 0.2.2) R package as described previously^61^. The binary filtered peak counts matrix from the Seurat object was used to build a Monocle3 Cell Data Set (cds) object. A Cicero cds object was created using the Cicero “make_cicero_cds” function with the parameter reduced_coordinates equal to the UMAP coordination in the Seurat object. Co-accessibility scores between peaks were calculated using the Cicero “run_cicero” function with the mouse mm10 genome. The gene activity score was calculated using the Cicero “build_gene_activity_matrix” function and normalized with the Cicero “normalize_gene_activities” function. The gene activity score was transformed with log(score*1000000 + 1). The unnormalized and normalized gene activity matrix were used to create a “RNA” assay in the Seurat object for downstream analysis. To find differentially gene activity between two groups of cells, the Seurat “FindMarkers” function was used.

### BMDM generation

Dissected femurs from 6-8 weeks old control and LysM^cre^KDM6B ^fl/fl^ mice were collected in ice cold complete RPMI 1640 media. Both epiphyses were removed before the bones were placed in sterile microfuge tubes and centrifuged at 500g for 5 minutes at 4 °C to extrude the bone marrow. The collected bone marrow was homogenized by pipetting followed by RBC lysis. The single cell suspension of bone marrow cells obtained, was counted in an automated Vicell cell counter before 2X10^6^ cells/well were cultured in Iscove’s Modified Dulbecco’s Medium (IMDM) supplemented with 10% FBS and 10ng/ml Macrophage colony-stimulating factor (M-CSF) (Biolegend), (growth media) in a 12 well plate. On Day 3, cells were resuspended in fresh growth media and on Day 7, cells were passaged in fresh complete IMDM containing 100ng/ml LPS (InvivoGen). On Day 8, the generated BMDMs were subjected to antigen presentation and phagocytosis assay as outlined below.

### ChIP sequencing

Both LPS stimulated and unstimulated BMDMs generated as described above, were subjected to ChIP using the MAGnify Chromatin Immunoprecipitation System (Applied Biosystems) according to the Manufacturer’s protocol. Briefly, following crosslinking with formaldehyde for 10 minutes, cells were resuspended in lysis buffer and subjected to sonication to shear the DNA to 150-300 kb fragments. For each immunoprecipitation reaction 10µg of anti- KDM6B (Active Motif) and anti H3K27me3 antibodies (Active Motif) were used. Following antibody incubation, samples were washed and the DNA was purified and eluted. DNA concentration was measured in Qubit using the dsDNA HS Assay kit (Invitrogen) and 10µg of DNA was sent for sequencing to the MD Anderson Cancer Center Advanced Technology Genomics Core where sequencing was done using the NextSeq500 instrument.

### ChIP sequencing data analysis

The quality of CHIPseq FASTQ sequences generated as described above, were assessed using FastQC, followed by mapping by bowtie2^62^ with mouse reference genome mm10. The bam files obtained from mapping were further processed using SAMBLASTER^63^ and SAMTOOLS^64^, for duplicate removal, sorting and indexing. SAMBAMBA^65^ was used to Normalize bam files per read counts by performing random sampling. The ChIP-seq signal enrichment over "Input" background was identified using Model based analysis of ChIP-seq (MACS) version 3^66^.The identified peaks were annotated using CHIPseeker^67^, clusterProfiler^68^ and AnnotationDbi. Quantitative comparisions of different datasets were performed with MAnorm^69^. The profile plots for specific genes were plotted using computeMatrix and plotProfile programs of deepTools^70^ and the gene tracks were plotted using EAseq version 1.111^71^.

### RNA isolation and real-time PCR

Total RNA was isolated from LPS stimulated BMDMs generated from control and LysM^cre^KDM6B^fl/fl^ mice using the TriZol (Invitrogen) method according to the Manufacturer’s protocol. 1µg of the RNA was reverse transcribed into complementary DNA (Superscript III cDNA kit from Invitrogen, USA) and the cDNA was used to measure the expression of genes of interest via Real Time PCR (Applied Biosystems 7500 Fast, USA). Primers used for real time PCR are as follows-MuSocs3 Forward- 5’-CGCCCAGGTCCTTTGCCTGA-3’ MuSocs3 Reverse- 5’-CCGCATCCCGGGGAGCTAGT-3’ MuMafb Forward- 5’-GGCAGGGAGTCTCTGTCGGC-3’ MuMafb Reverse- 5’-CAGGCCCTCCGACCCCATCT-3’

### Phagocytosis assay with beads

Uncoated carboxylate modified polystyrene fluorescent orange beads (Sigma Aldrich) were added to 1X10^5^ BMDMs generated using the protocol outlined above, at a ratio of 500:1. The cells were incubated at 37 °C and 5% CO_2_ for 4 hours before being washed in PBS, fixed in 1% PFA and acquired in a flow cytometer to monitor uptake of beads by the cells.

### Phagocytosis assay with GL261 cells

CellTrace Far Red (Invitrogen) labelled GL261 cells were mixed with CTV labelled BMDMs (generated as described above) at at ratio of 1:1 and incubated at 37°C and 5% CO_2_ for 2 hours before the mixture was washed with PBS, fixed in 1% PFA and acquired in a flow cytometer to measure the uptake of GL261 cells by the BMDMs.

### Antigen presentation assay

BMDMs generated above were pulsed with GP100 (1µg/ml) or LCMV (1µg/ml, negative control) peptides (AnaSpec). Additionally, naïve CD8 T cells were isolated from the spleen of pmel mice (having CD8 T cells bearing T cell receptors specific for the gp100 antigen) by magnet-assisted cell sorting (Naïve CD8+ T cell isolation kit, Miltenyi Biotec) and stained with Cell Trace Violet (CTV, Invitrogen). Subsequently, the antigen pulsed macrophages and stained naïve CD8 T cells were co-cultured at a ratio of 1:2 for 3 days, washed, fixed and subjected to flow cytometry (BD LSR II) to measure CTV dilution as a measure of T cell proliferation in response to antigen presentation by the different BMDMs. FlowJo software v10 was used for analysis.

### GSK-J4 and anti-PD-1 treatment regimen

5–7-weeks old C57BL/6 mice bearing GL261 or CT-2A tumors were treated with KDM6B inhibitor (GSK-J4; Sigma-Aldrich); 1mg per mouse, in 200ul volume (3% DMSO+ 97% sterile water - vehicle) via oral gavage daily starting from day 3 post tumor inoculation to the end of the experiment. GL261 tumor bearing micewere also injected intraperitoneally with 200µg, 100µg and 100µg of α-PD1 (RMP1-14; Bio X Cell) on day 7, 10 and 13 respectively and sacrificed on day 17 for downstream analysis.

### Mass cytometry to study TME

GL261 and CT-2A tumor tissues were dissected, processed, stained with antibodies and acquired in a mass cytometer as previously described. Surface and intracellular antibodies used for this mass cytometry are mentioned in Supplementary Table 2.

### Mass cytometry analysis

Files were manually gated in FlowJo v10 by using iridium for cells, event length for singlets, cisplatin for live/dead discrimination and using CD45 lineage marker for immunecells. Clustering analysis was performed using the FlowSOM and ConsensusClusterPlus packages as previously described ^72^

### Statistical analyses

R v4.0.2 and GraphPad Prism software v9 was used for the statistical analyses. The individual tests performed have been indicated in the figure legends. All in-vivo experiments had two to three independent replicates.

### Data availability

Raw reads for the single cell RNA sequencing, single cell ATAC sequencing and ChIP-sequencing will be deposited in European Genome-phenome Archive (EGA) which will be available upon acceptance of the manuscript.

### Code availability

The scRNA seq, scATAC seq and CHIP seq analyses presented in the manuscript has been performed with open source algorithms as described in the method section. Further details will be made available by authors upon request.

## Extended Data figure legend

**Extended Data Fig. 1: (A)** Dotplot showing the average expression of indicated genes as well as the percentage of cells expressing the gene in the indicated *CD45*+ immune cell clusters in the TME of patients with GBM (n=5) shown in Fig. 1A. **(B)** UMAP plot of scRNA seq data depicting the expression of *KDM6B* gene in the different immune cell subsets (*CD45*+ cells). **(C)** Violin plots demonstrating the expression level of indicated genes in the different myeloid cell clusters from Fig. 1B.

**Extended Data Fig. 2: (A)** UMAP plot of scRNA seq data showing *CD45*+ immune cell subsets in the GBM TME derived from patients with GBM (n=4)^32^. (**B)** Violin plots demonstrating the expression level of indicated genes in the different immune cell clusters. **(C)** UMAP plot of scRNA seq data depicting *CD45*+ immune cell subsets in the GBM TME derived from patients with GBM (n=20)^37^.**(D)** Violin plots representing the expression level of indicated genes in the different immune cell clusters.

**Extended Data Fig. 3: (A)** Hematoxylin and Eosin stained GBM tumor sections. Each section represent one patient. Scale included in the images. **(B)** Representative figures showing immunohistochemical staining for KDM6B in GBM tissue samples from 3 patients (patient number: 6, 2 and 3, Supplemental Table 1)

**Extended Data Fig. 4: (A, B)** UMAP plots of matched scRNA seq data from patient 2 and patient 3 (Supplementary Table 1) showing *CD45*+ immunecell subsets in the GBM TME used to embed single cells to their spatial coordinates in tissue sections by applying CellTrek. **(C, D)** Gene expression data of all the different immune cell clusters from matched patients plotted on spatial coordinates. **(E, F)**Gene expression data from the *Kdm6b* expressing myeloid cell clusters (black) and T cell clusters (yellow) plotted on spatial coordinates of two matched patients.

**Extended Data Fig.5 (A-C)** Dotplots showing the average gene activity score of genes of interest as well as percentage of cells in the cluster expressing the gene in the indicated clusters **(A)** shown in Fig. 1F, and **(B,C)** shown in Fig. S4A,B.

**Extended Data Fig.6: (A)** Schematic representation demonstrating generation of the LysM^cre^KDM6B^fl/fl^ genetic murine model. **(B)** Representative image of an agarose gel showing bands depicting PCR amplified DNA from *Kdm6b* deleted homozygous mice (single 400bp band), *Kdm6b* deleted heterozygous mice (both 368 and 400bp bands), and control homozygous mice (single 368bp band).**(C)** t-SNE plots and box and whisker plots depicting the identity and abundance of different immune cell populations present in the indicated anatomical locations in control and LysM^cre^KDM6B^fl/fl^ mice as determined from CyTOF analysis (n=3/group).

**Extended Data Fig. 7: (A)** Heatmap showing the expression of genes of interest in the different *Cd45+* immune cell clusters (shown in Fig. 2C). **(B)** Bar graphs depicting the frequencies of the different intratumoral immune cell subsets in control and LysM^cre^KDM6B^fl/fl^ mice. **(C)** Bar graph representing the ratio of intratumoral CTLs and Tregs in control and LysM^cre^KDM6B^fl/fl^ mice as determined from scRNA seq. **(D)** Violin plots depicting the expression level of *Lyz2* in different immune cell clusters in control and LysM^cre^KDM6B^fl/fl^ mice. Data representative of two independent scRNA seq experiments.

**Extended Data Fig. 8: (A)** t-SNE plot of CyTOF data demonstrating different immunecell subsets (CD45+ cells) in the GBM (CT-2A) TME of control and LysM^cre^KDM6B^fl/fl^ mice (n=10/group). **(B)** Heatmap showing the expression of protein markers of interest in the indicated immune cell clusters as determined by mass cytometry. **(C)** Box and whisker plots representing the relative frequencies of indicated immune cell clusters from control and LysM^cre^KDM6B^fl/fl^ mice as determined by CyTOF (control n=10/group, LysM^cre^KDM6B^fl/fl^ n=8/group). Two-tailed Student’s t-test was performed (*p<0.05, **p<0.01, ****p<0.0001). **(D)** Bar plots depicting the ratio of intratumoral CTLs and Tregs in control and LysM^cre^KDM6B^fl/fl^ mice as determined by CyTOF.

**Extended Data Fig. 9: (A)** Heatmap showing the expression of genes of interest in the different myeloid cell clusters (shown in Fig. 3A-right panel). **(B)** Bar plots representing the frequencies of intratumoral myeloid clusters from control and LysM^cre^KDM6B^fl/fl^ mice. Data representative of two independent experiments.

**Extended Data Fig. 10: (A)** UMAP demonstrating the CTL and Treg clusters in the GBM (GL261) TME of control and LysM^cre^KDM6B^fl/fl^ mice determined by scATAC seq (as shown in Fig. 4 A-B). Bar graphs depicting the frequencies and ratio of intratumoral CTLs and Tregs in control and LysM^cre^KDM6B^fl/fl^ mice as determined from scATAC seq. **(B)** Coverage plots depicting the chromatin accessibility of the indicated genes in the CTL and Treg clusters.

**Extended Data Fig. 11: (A)** Heatmap showing the expression of genes of interest in the indicated myeloid cell clusters (shown in Fig. 4D). **(B, C)** Coverage plots depicting accessibility of indicated chromatin regions (peaks) in genes of interest. **(D)** Representative gating strategy on FlowJo for analysis of flow cytometry data showing GL261 phagocytosis by BMDMs (shown in Fig. 5J). **(E)** Representative pseudocolor flow cytometry plots showing the percentage of proliferated CD8 T-cells (gated, CTV negative), upon co-culture with LCMV pulsed control and *Kdm6b* deficient BMDMs.

**Extended Data Fig.12:** Heatmap representing the expression of genes of interest in the indicated *Cd45*+ immune cell clusters as determined by scRNA seq (shown in Fig. 6C).

**Extended Data Fig. 13: (A)** Representative axial MRI images of CT-2A tumor from vehicle treated mice (left panel) and GSK-J4 treated mice (right panel), taken on day 14 post tumor inoculation. **(B)** Box and whisker plot showing the difference in CT-2A tumor volumes (determined from MRI) between vehicle and GSK-J4 treated mice (n=10/ group). Two-tailed Student’s t-test was performed (*p<0.05). **(C)** Box and whisker plot depicting the difference in CT-2A tumor weight (harvested on day 22 post tumor inoculation) between vehicle and GSK-J4 treated mice (n=10/group). Two-tailed Student’s t-test was performed (*p<0.05). **(D)** Heatmap demonstrating the expression of protein markers of interest in the indicated CD45+ immune cell clusters in the GBM (CT-2A) TME of vehicle & GSK-J4 treated mice as determined by mass cytometry.**(E)** t-SNE plot of CyTOF data depicting different immune cell subsets (CD45+ cells) in the GBM (CT- 2A) TME of vehicle and GSK-J4 treated mice (n=5/group). **(F)** Box and whisker plots representing the relative frequencies of indicated immune cell clusters from vehicle and GSK-J4 treated mice as determined by CyTOF (n=5/group). Two-tailed Student’s t-test was performed (**p<0.01).

**Extended Data Fig. 14:** Heatmap showing the expression of protein markers of interest in the indicated CD45+ immune cell clusters as determined by mass cytometry (shown in Fig. 6H).

## Acknowledgment

This research is supported by the MD Anderson Physician Scientist Award (S.G), Khalifa Physician Scientist Award (S.G), Andrew Sabin Family Foundation Fellows Award (S.G) and Clinic and Laboratory Integration Program Award (S.G). We acknowledge Liangwen Xiong, Baoxiang Guan, Derek Ng Tang, Sydney Kemp and Alison Jung for technical assistance. We acknowledge the CATALYST-working group at MD Anderson Cancer Center for human GBM tumor samples. CATALYST is supported by the MD Anderson GBM Moon Shot™. Dr. Sharma is a member of the Parker Institute for Cancer Immunotherapy.

## Author information

S.G. developed the project, designed the experiments, analyzed data, wrote the manuscript and acquired funding. D.R, S.M.N, P.S performed the experiments, analyzed data and wrote the manuscript. Y.C, performed bioinformatics analyses. J.Z, M.H, S.A helped with the murine experiments. B.P.K, C.P and F.L provided human GBM tumor samples. M.M and S.J performed the H&E, IHC and IF staining of human GBM samples. S.B and Z.H helped with the human scRNA-sequencing and VISIUM analysis. P.S oversaw the study, provided scientific input, edited the manuscript and acquired funding.

