## Supplementary Information for "Myeloid-specific KDM6B inhibition sensitizes Glioblastoma to PD1 blockade"

#### Extended Data Fig. 1

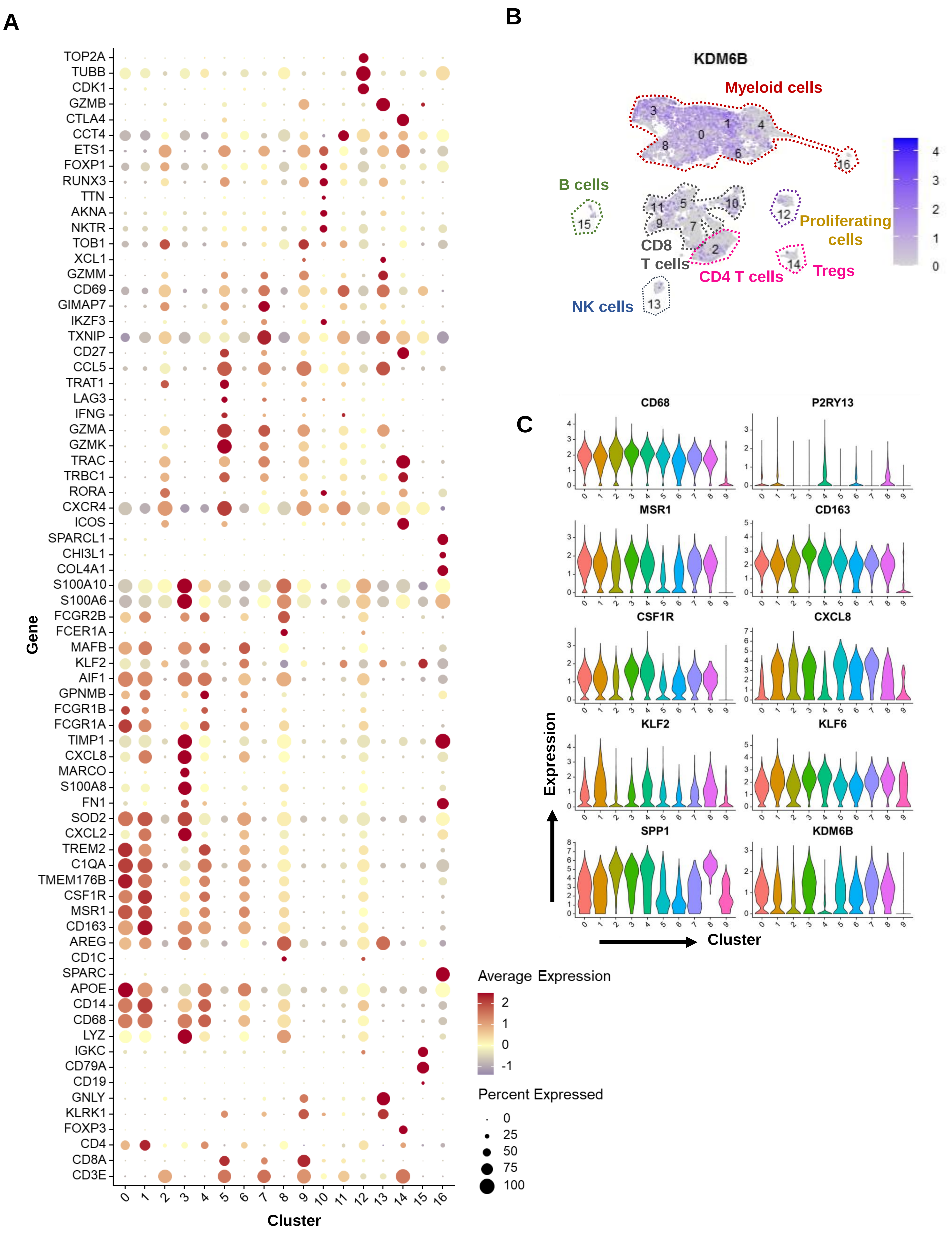

#### Extended Data Fig. 2

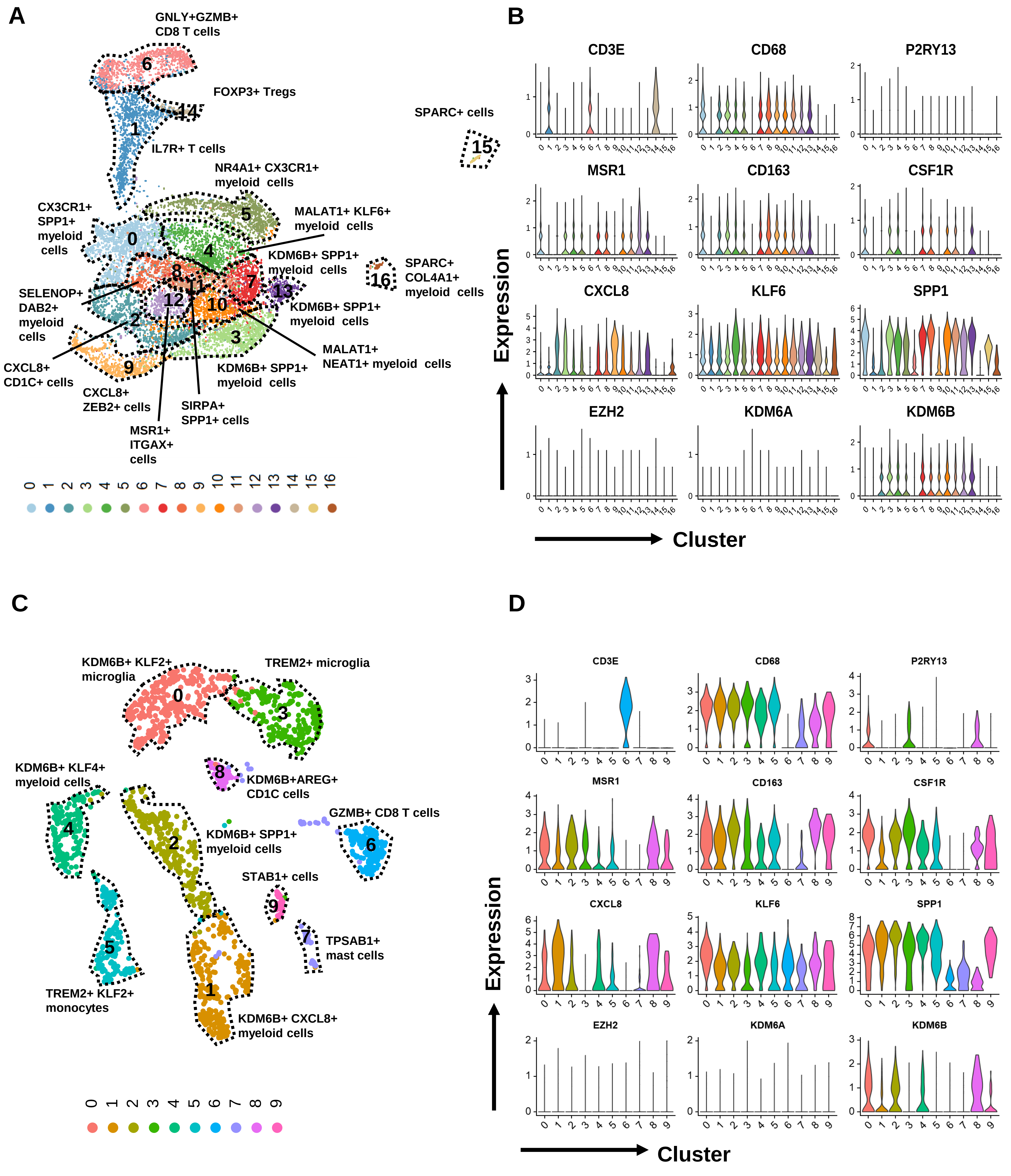

**Extended Data Fig. 3**

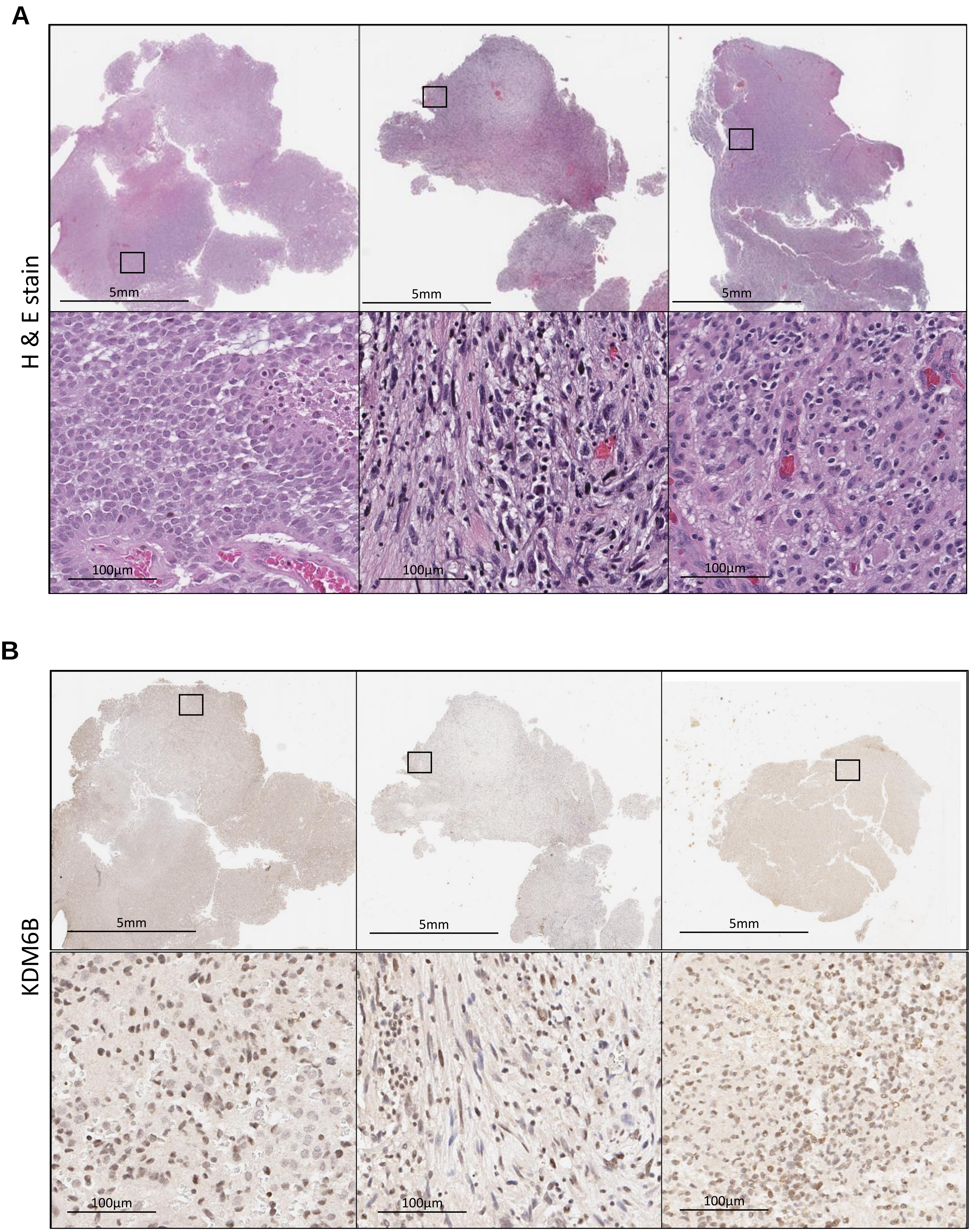

#### Extended Data Fig. 4

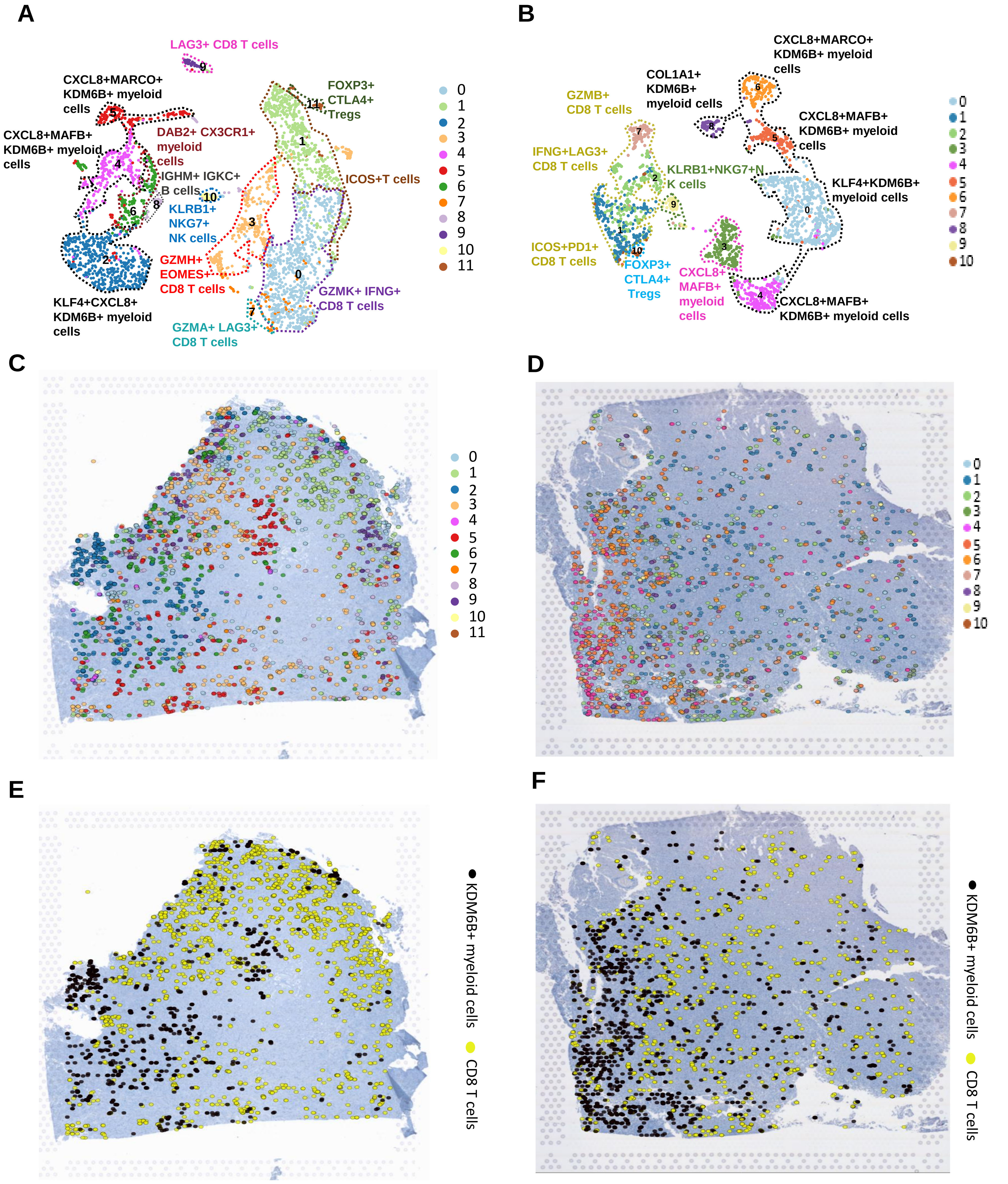

**Extended Data Fig. 5**

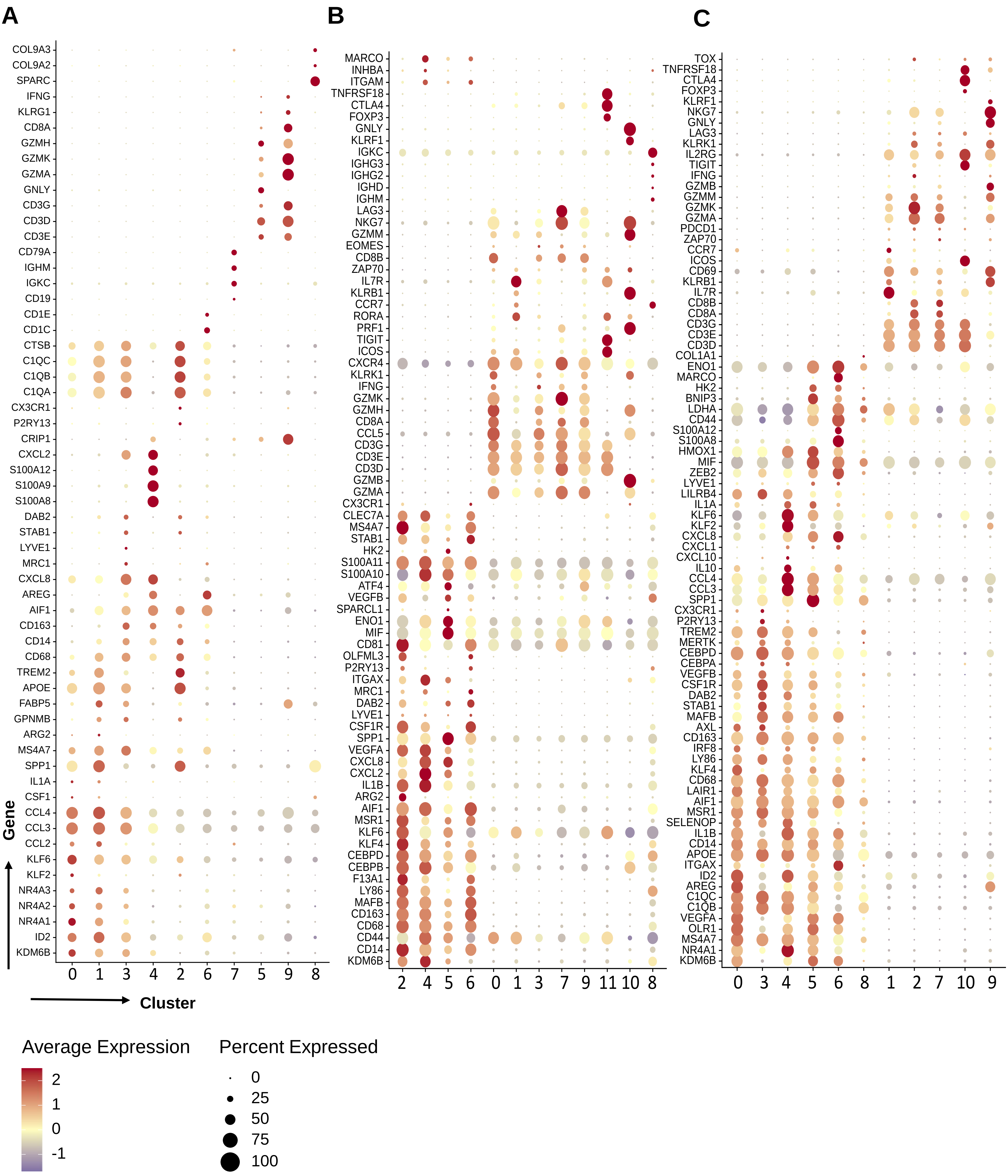

**Extended Data Fig. 6**

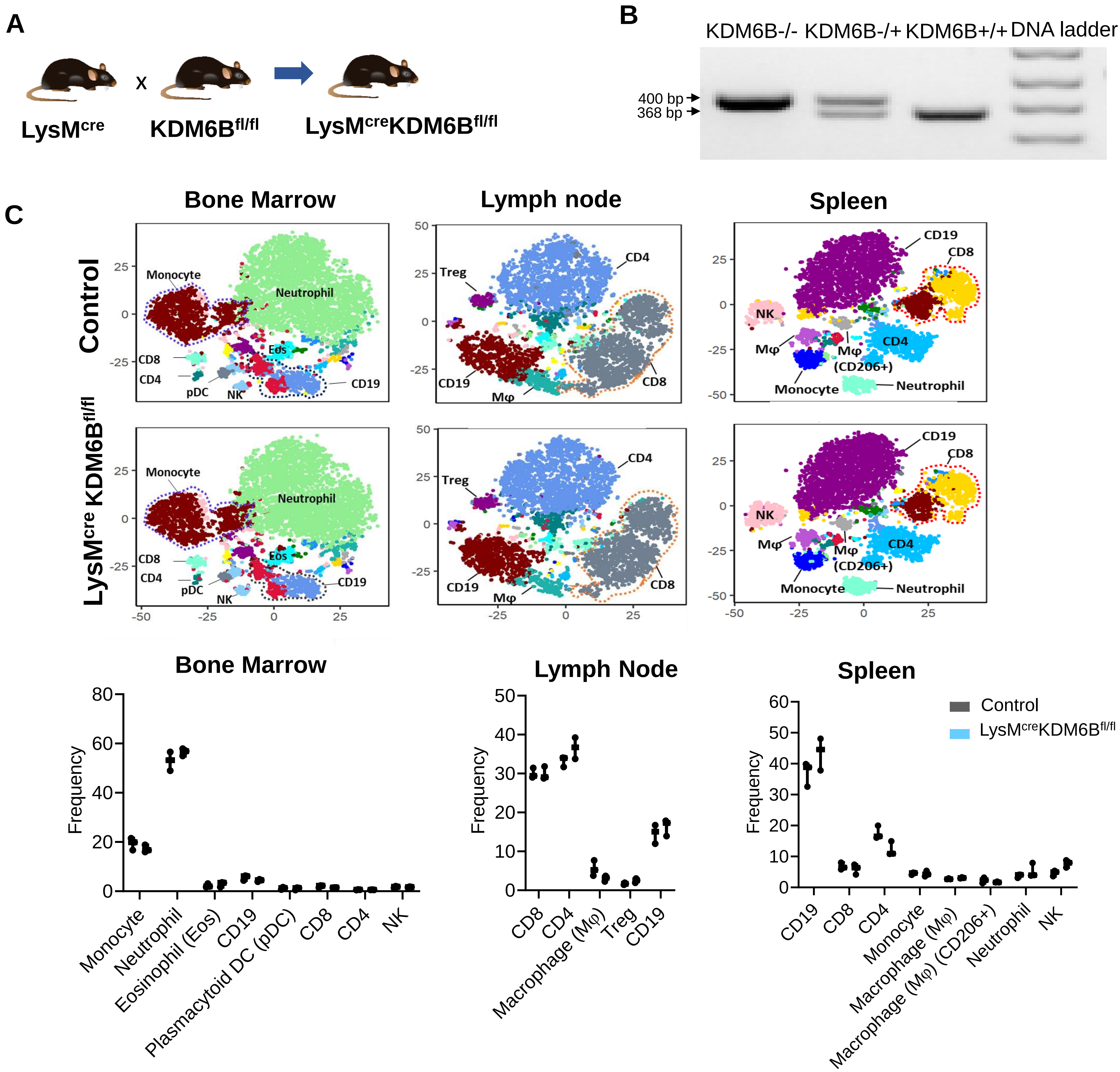

**Extended Data Fig. 7**

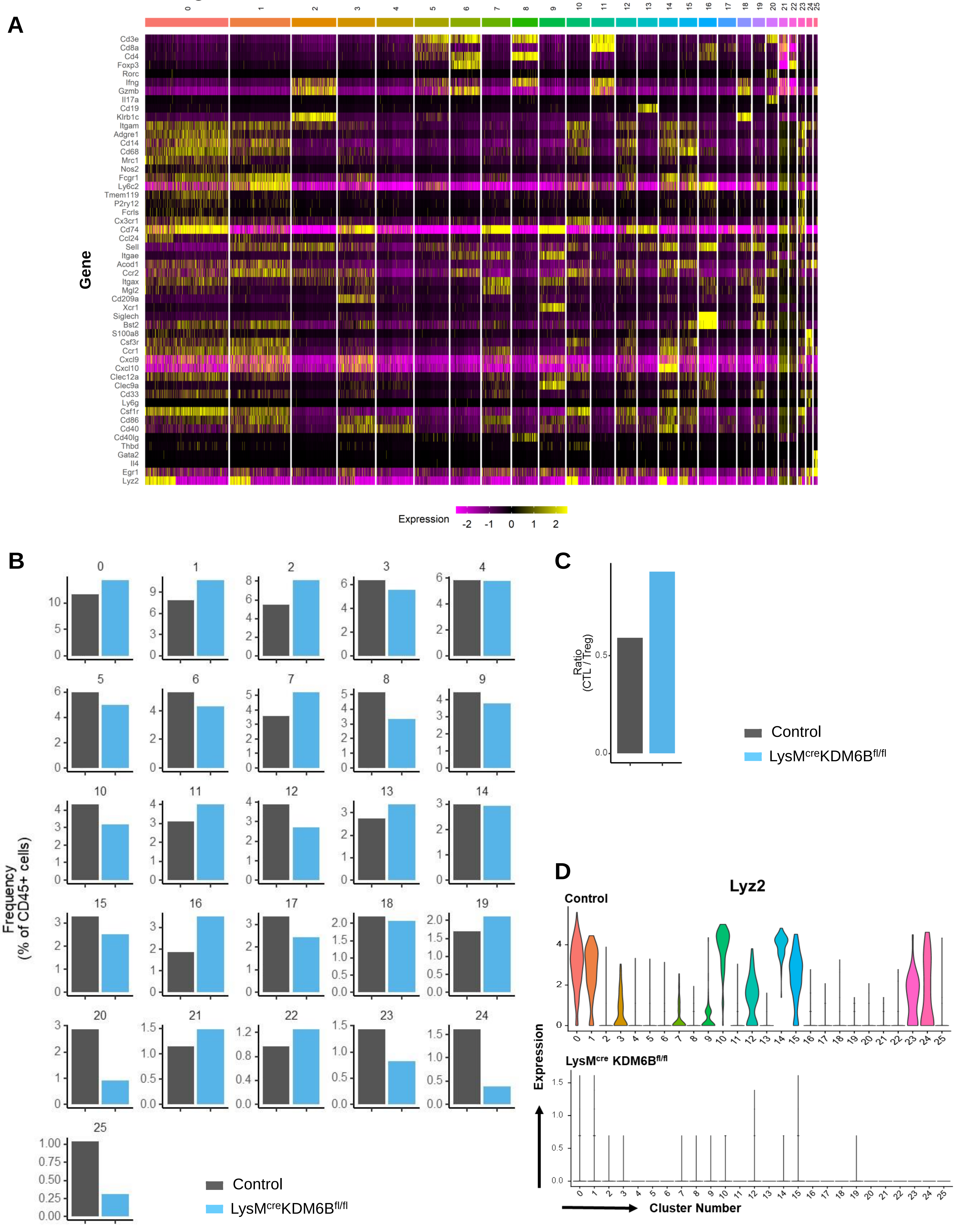

### Extended Data Fig. 8

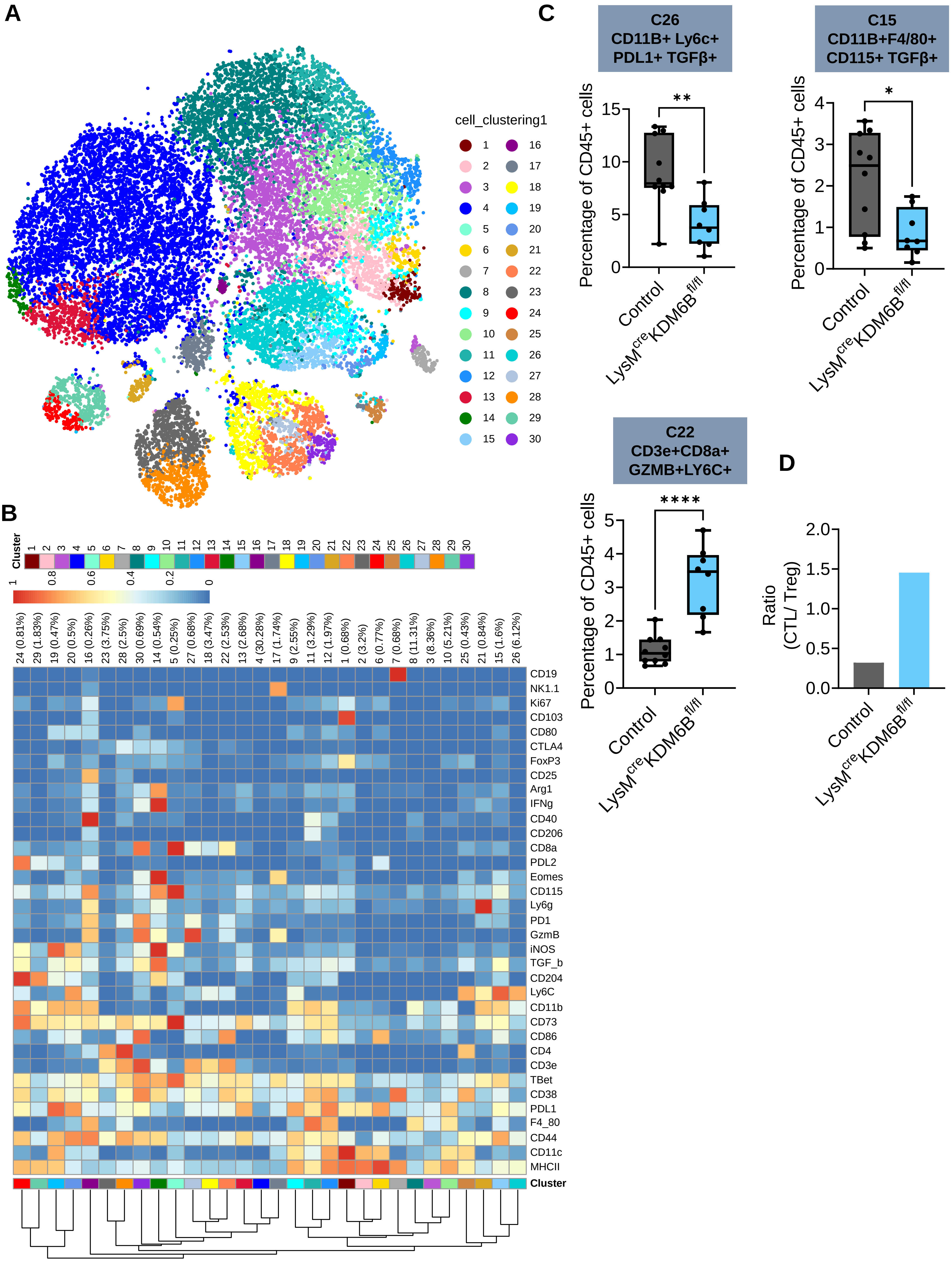

**Extended Data Fig. 9**

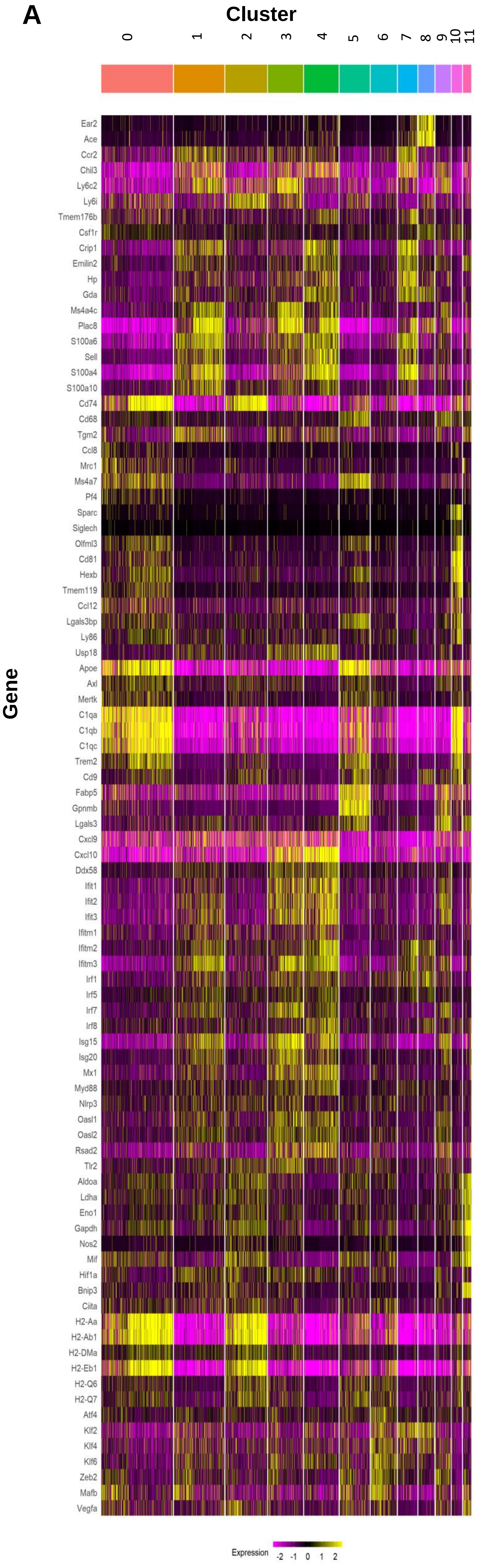


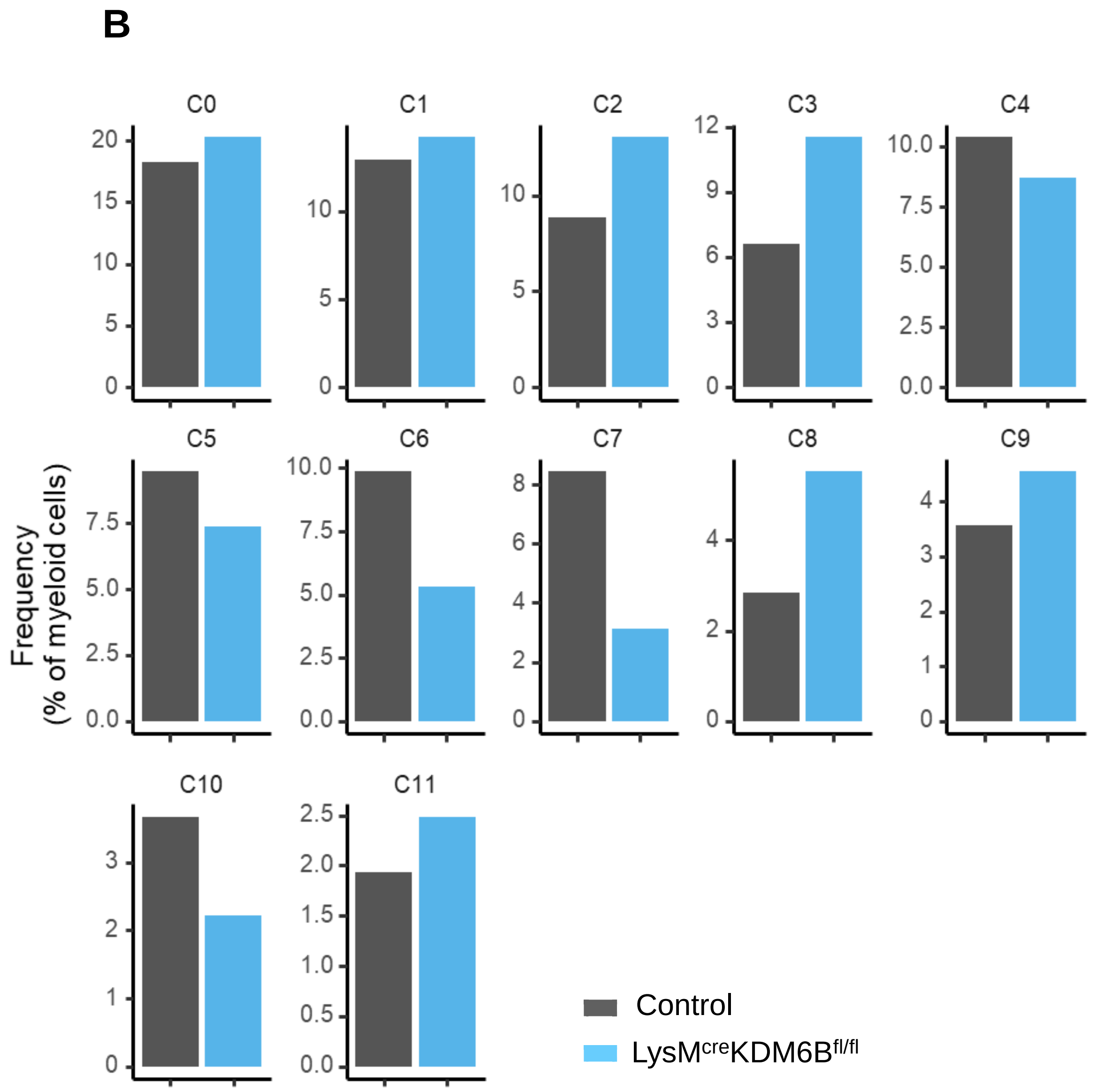

### Extended Data Fig. 10

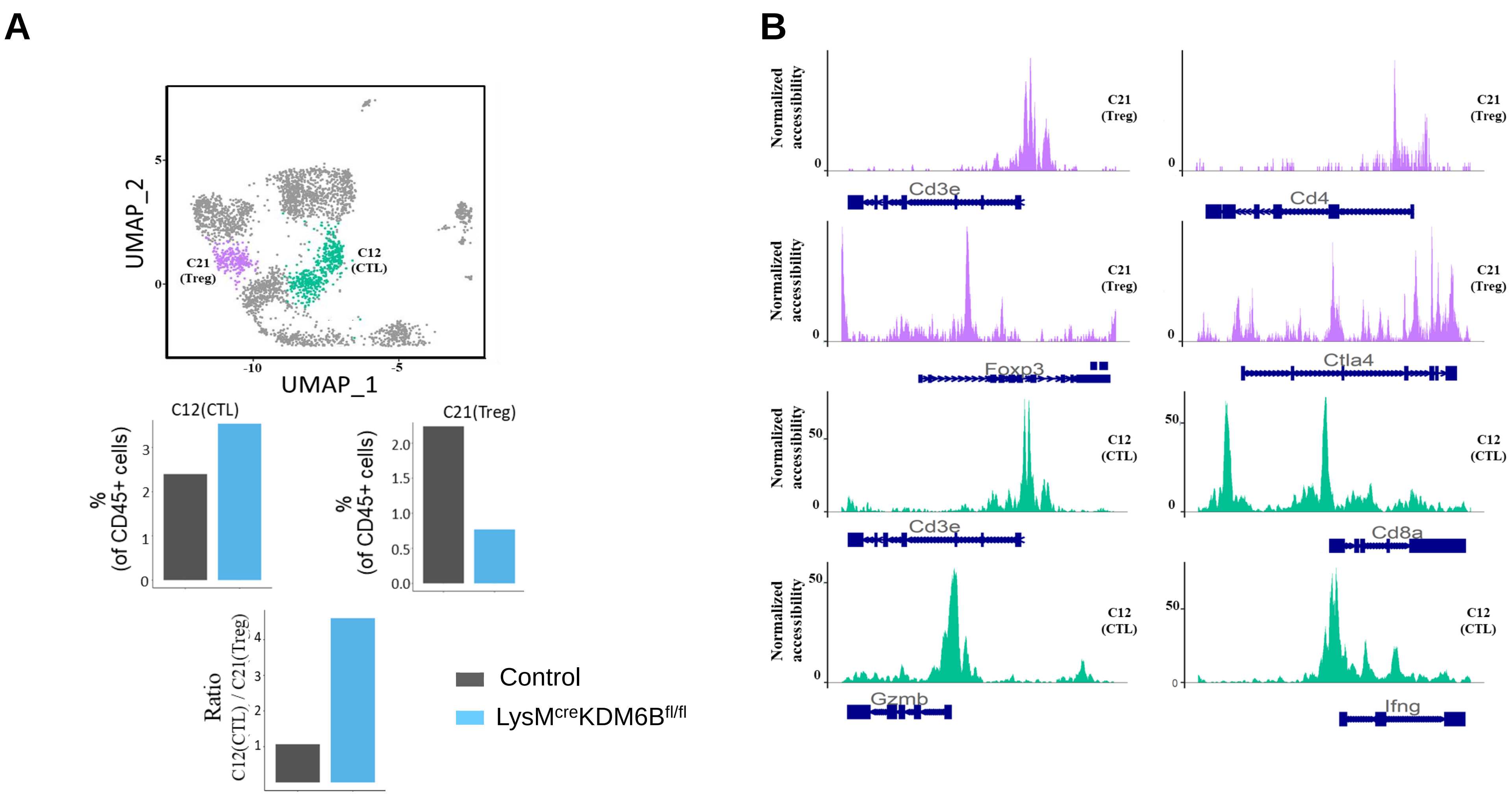

#### Extended Data Fig. 11

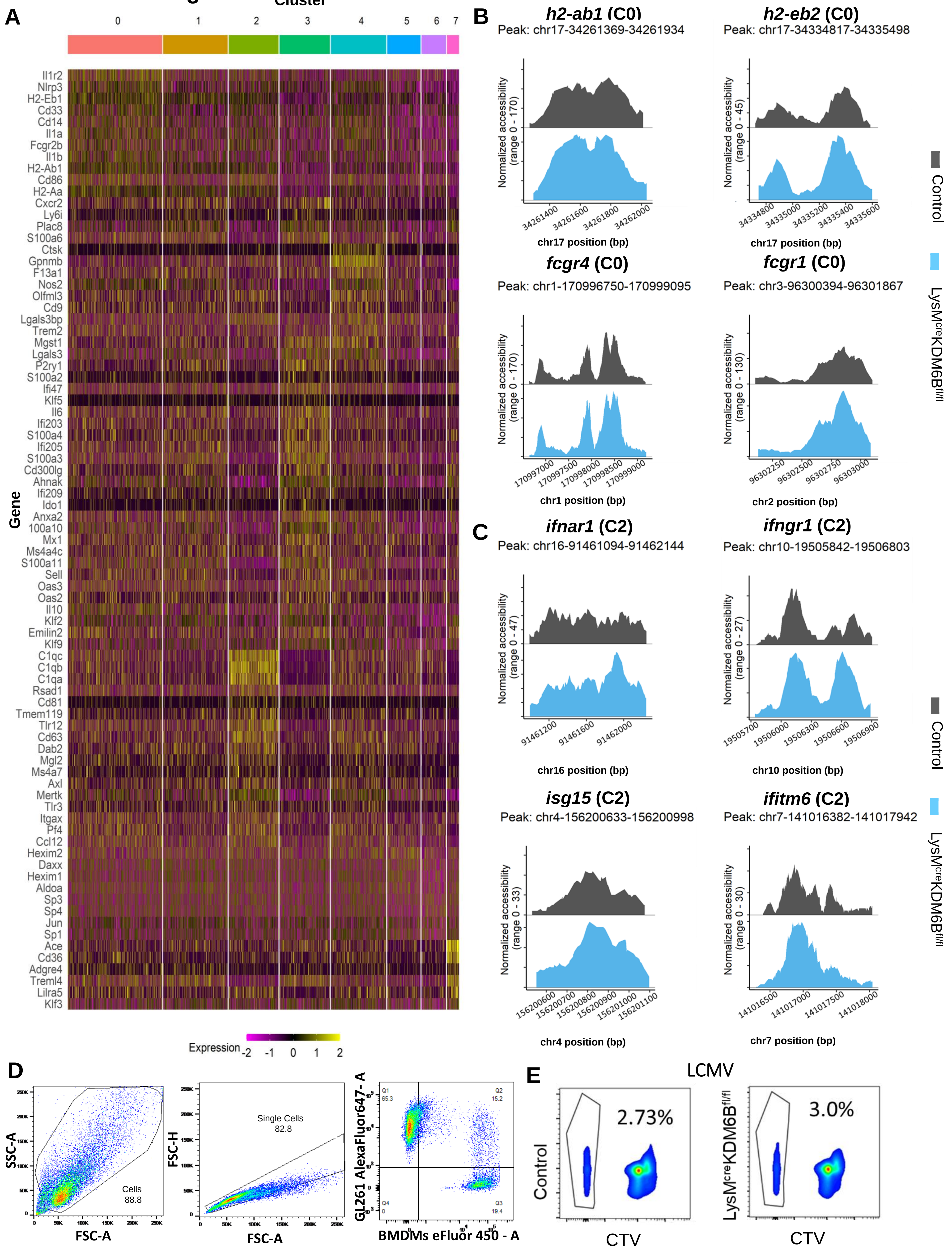

Extended Data Fig. 12

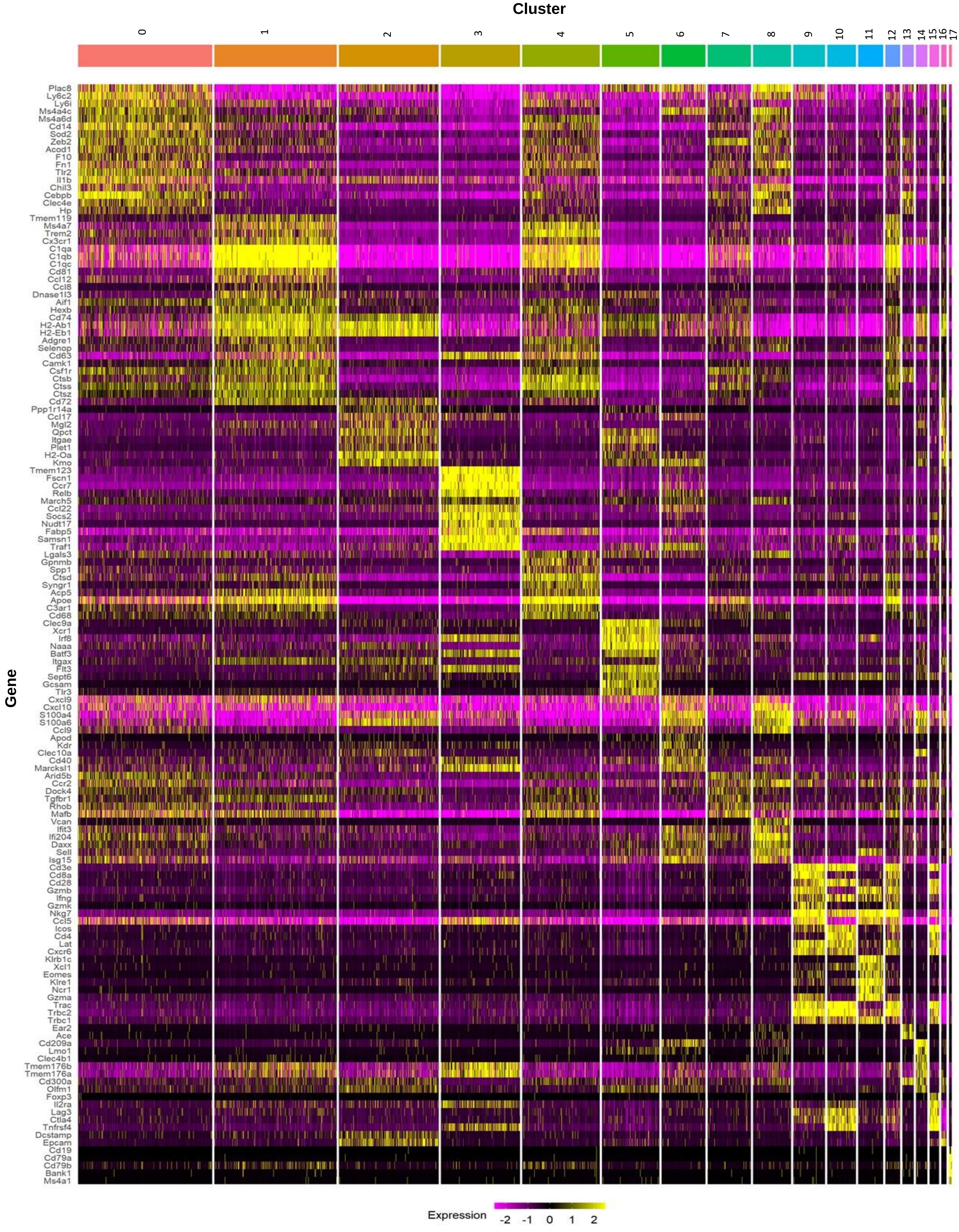

#### Extended Data Fig. 13

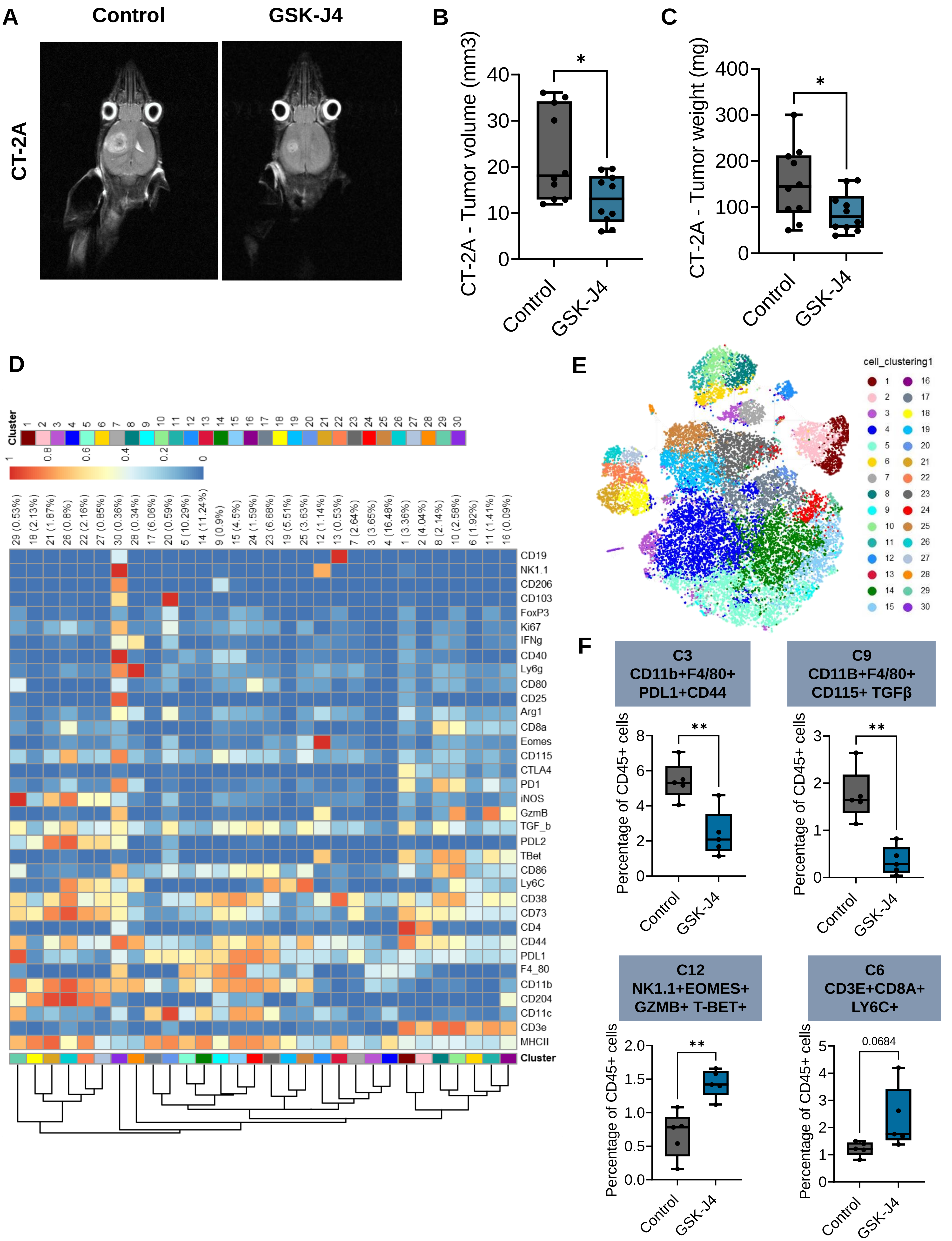

**Extended Data Fig. 14**

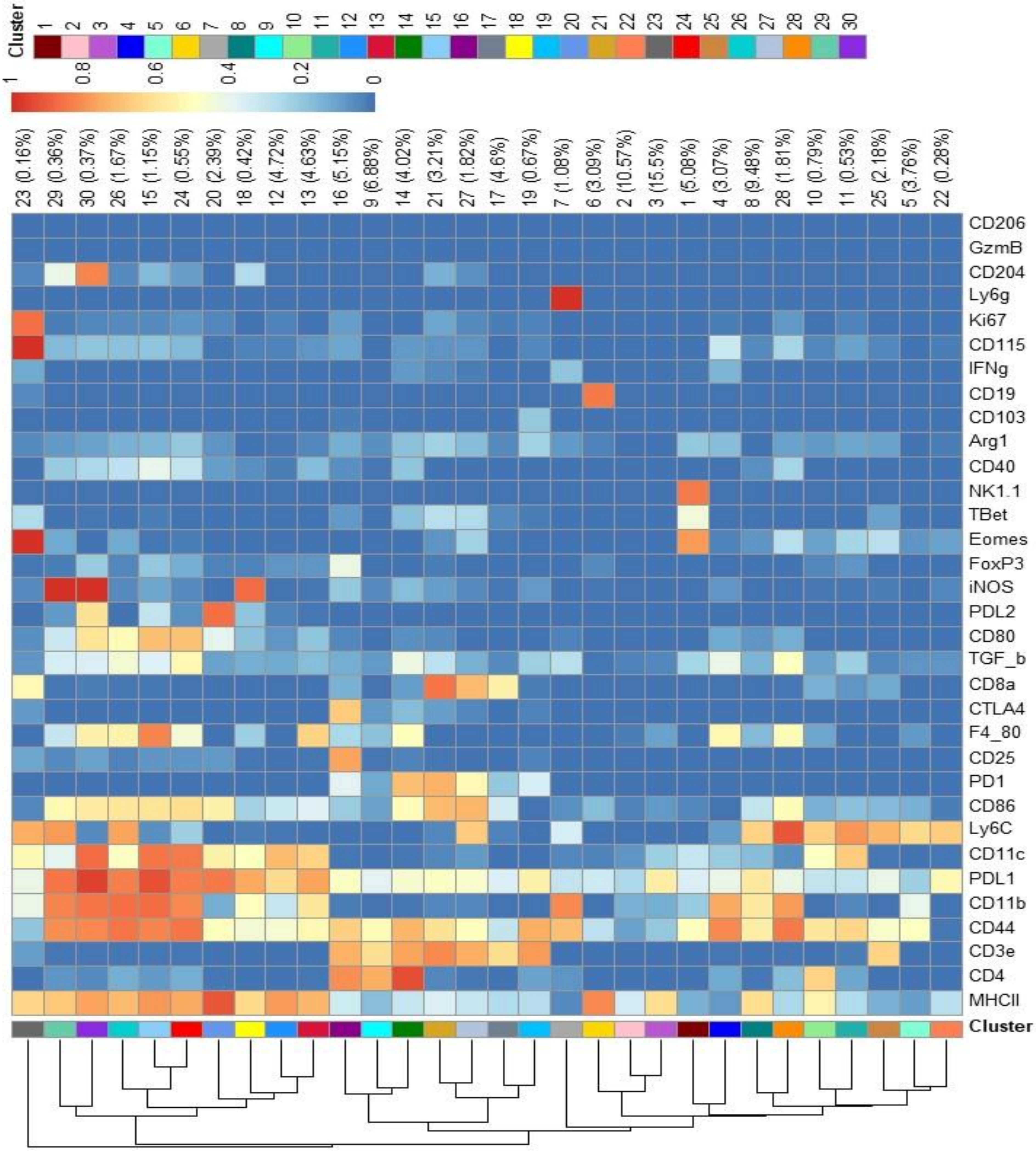

### Supplementary Table 1: Patient characteristics

| Patient # | 1 | 2 | 3 | 4 | 5 | 6 |
|---|---|---|---|---|---|---|
| Previous chemoradiation | No | No | No | No | No | No |
| Previous surgery | Yes | Yes | No | No | No | No |
| Previous type of surgery (biopsy, subtotal resection, gross total resection) | Biopsy | Subtotal resection | NA | NA | NA | NA |
| On steroids day prior to surgery | No | Yes | Yes | Yes | Yes | No |
| Steroid dose prior to surgery | NA | 2mg PO daily | 4mg PO TID | 4mg PO Q6H | 2mg PO BID | NA |
| Received a perioperative dose of steroids | Yes | Yes | Yes | Yes | Yes | Yes |
| Perioperative dose of steroids | 10mg IV once | 6mg IV Q6H | 10mg IV once | 10mg IV once | 10mg IV once | 10mg IV once |
| Pathology type | Glioblastoma | Glioblastoma | Glioblastoma | Glioblastoma | Glioblastoma | Glioblastoma |
| Pathology WHO grade | 4 | 4 | 4 | 4 | 4 | 4 |
| IDH mutant | No | No | No | No | No | No |
| 1P/19Q codeleted | No | No | No | No | No | No |
| Procedure | Craniotomy resection | Craniotomy Resection | Craniotomy Resection, Awake | Craniotomy resection | Craniotomy resection | Craniotomy resection |
| Assays performed | Single cell RNA sequencing | Single cell RNA sequencing, Visium | Single cell RNA sequencing, Visium | Single cell RNA sequencing | Single cell RNA sequencing | Single cell RNA sequencing, Visium |

### Supplementary Table 2: Key Resource table with list of antibodies

#### CyTOF antibodies

| Marker | Clone | Company | Catalog number | Dilution |
|---|---|---|---|---|
| Anti-CD45 | 30-F11 | Fluidigm | Cat # 3089005B | 1:200 |
| Anti-MHCII | M5/114.15.2 | Biolegend | Cat # 107637 | 1:800 |
| Anti-CD11c | N418 | Fluidigm | Cat # 3142003B | 1:50 |
| Anti-CD4 | RM4-5 | Biolegend | Cat # 100506 | 1:200 |
| Anti-CD115 | AFS98 | Fluidigm | Cat # 3144012B | 1:50 |
| Anti-CD73 | AD2 | Biolegend | Cat # 344002 | 1:400 |
| Anti-CD8a | 53-6.7 | Fluidigm | Cat # 3146003B | 1:400 |
| Anti-Ly6g | 1A8 | Biolegend | Cat # 127637 | 1:200 |
| Anti-CD11b | M1/70 | Biolegend | Cat # 101249 | 1:1600 |
| Anti-CD19 | 6D5 | Fluidigm | Cat # 3149002B | 1:200 |
| Anti-Ly6C | HK1.4 | Biolegend | Cat # 128039 | 1:400 |
| Anti-CD25 | 3C7 | Fluidigm | Cat # 3151007B | 1:50 |
| Anti-CD3ε | 145-2C11 | Fluidigm | Cat # 3152004B | 1:100 |
| Anti-PD-L1 | 10F.9G2 | Fluidigm | Cat # 3153016B | 1:200 |
| Anti-CTLA-4 | UC10-4B9 | Fluidigm | Cat # 3154008B | 1:100 |
| Anti-TBET | 4B10 | Biolegend | Cat # 644825 | 1:150 |
| Anti-GzmB | QA16A02 | Biolegend | Cat # 372202 | 1:100 |
| Anti-Foxp3 | FJK-16s | Fluidigm | Cat # 3158003A | 1:100 |
| Anti-PD-1 | J43 | Fluidigm | Cat # 3159023B | 1:100 |
| Anti-CD38 | 90 | Biolegend | Cat # 102702 | 1:400 |
| Anti-iNOS | CXNFT | Fluidigm | Cat # 3161011B | 1:100 |
| Anti-CD204 | 7G5C33 | Biolegend | Cat # 676202 | 1:100 |
| Anti-PD-L2 | Ty25 | Biolegend | Cat # 107202 | 1:100 |
| Anti-TGF-b | TW7-16B4 | Fluidigm | Cat # 3164014B | 1:200 |
| Anti-IFNg | XMG1.2 | Biolegend | Cat # 505802 | 1:100 |
| Anti-Eomes | Dan11mag | Invitrogen/eBioscience | Cat # 14-4875-82 | 1:100 |
| Anti-CD80 | 16-10A1 | Biolegend | Cat # 104710 | 1:800 |
| Anti-Arg1 | 8C9 | Santa Cruz Biotechnology | Cat # sc-47715 | 1:300 |
| Anti-CD206 | C068C2 | Biolegend | Cat # 141702 | 1:200 |
| Anti-NK1.1 | PK136 | Fluidigm | Cat # 3170002B | 1:100 |
| Anti-CD44 | IM7 | Fluidigm | Cat # 3171003B | 1:400 |
| Anti-GATA3 | TWAJ | Invitrogen/eBioscience | Cat # 14-9966-82 | 1:100 |
| Anti-SIRP1a | P84 | Biolegend | Cat # 144035 | 1:400 |
| Anti-CD40 | 23-Mar | Biolegend | Cat # 124602 | 1:200 |
| Anti-F4/80 | BM8 | Invitrogen/eBioscience | Cat # 14-4801-82 | 1:100 |
| Anti-CD86 | GL-1 | Biolegend | Cat # 105002 | 1:100 |
| Anti-CD62L | MEL-14 | Biolegend | Cat # 104402 | 1:100 |
| Anti-SiglecH | 551 | Biolegend | Cat # 129602 | 1:100 |
| Anti-SiglecF | E50-2240 | BDBiosciences | Cat # 552125 | 1:100 |
| Anti-CD117 | c-kit | Invitrogen/eBioscience | Cat # 14-1172-82 | 1:200 |
| Anti-fceR | MAR-1 | Biolegend | Cat # 134302 | 1:100 |
| Anti-ICOS | ISA-3 | eBioscience | Cat # 14-9948-82 | 1:100 |
| Anti-LAG3 | `874501 | R&D | Cat # MAB23193SP | 1:100 |
| Anti-CD103 | 2 e7 | Biolegend | Cat # 121402 | 1:100 |
| Ir DNA-Intercalator | | Fluidigm | Cat # 201192A | |
| cisplatin | | Fluidigm | Cat# 201064 | |

#### ChIP antibodies

| Marker | Clone | Company | Catalog number | Dilution |
|---|---|---|---|---|
| Anti-KDM6B | 67-A2 | Active Motif | Cat#61387 | 10ug per pull down reaction |
| H3K27ME3 | | Active Motif | Cat#39155 | 10ug per pull down reaction |

#### Flow antibodies

| Marker | Clone | Company | Catalog number | Dilution |
|---|---|---|---|---|
| Anti-CD45 Pacific Blue | 30-F11 | Biolegend | Cat # 103126 | 1:200 |
| Anti-CD3E FITC | 17A2 | eBioscience | Cat # 11-0032-82 | 1:200 |
| Anti-CD11B APC | M1/70 | Biolegend | Cat # 101212 | 1:200 |
| Live/ Dead Pacific Orange | | Invitrogen | Cat # L34968 | 1:500 |

#### Immunofluroscence antibodies

| Marker | Clone | Company | Catalog number | Dilution |
|---|---|---|---|---|
| Anti-CD3E | | Agilent | Cat # A0452 | 1:200 |
| Anti-CD8 | C8/144b | Abcam | Cat # ab17147 | 1:100 |
| Anti-CD68 | PGM-1 | Agilent | Cat # M0876 | 1:25 |
| Anti-CD163 | 10D6 | Abcam | Cat # ab201461 | 1:20 |

#### Antibodies for Immunohistochemistry

| Marker | Clone | Company | Catalog number | Dilution |
|---|---|---|---|---|
| Anti-KDM6B | | Invitrogen | Cat # PA5-32192 | 1:200 |
